# Identifying causal genotype-phenotype relationships for population-sampled parent-child trios

**DOI:** 10.1101/2024.12.10.627752

**Authors:** Yushi Tang, Irineo Cabreros, John D. Storey

## Abstract

The process by which genes are transmitted from parent to child provides a source of randomization preceding all other factors that may causally influence any particular child phenotype. Because of this, it is natural to consider genetic transmission as a source of experimental randomization. In this work, we show how parent-child trio data can be leveraged to identify causal genetic loci by modeling the randomization during genetic transmission. We develop a new test, the transmission mean test (TMT), together with its unbiased estimator of the average causal effect, and derive its causal properties within the potential outcomes framework. We also prove that the transmission disequilibrium test (TDT) is a test of causality as a complementary case of the TMT for the affected-only design. The TMT and the TDT differ in the types of traits that they can handle and the study designs for which they are appropriate. The TMT handles arbitrarily distributed traits and is appropriate when trios are randomly sampled; the TDT handles dichotomous traits and is appropriate when sampling is based on a child’s trait status. We compare the transmission-based methods with established approaches for genotype-phenotype analyses to clarify conditions appropriate for each method, what conclusions can be drawn by each one, and how these methods can be used together.

## 1 Introduction

The ultimate goal of understanding the genotype-phenotype relationship is to uncover genetic variants that are causal for the phenotype. The gold standard statistical framework for establishing causality is the potential outcomes framework [1–3]. This paper investigates how human trio data (two parents and their child) can enable tests of causality based on comparisons of trait level potential outcomes corresponding to different alleles transmitted from the parents to the child. The motivation lies in the connection between the meiotic process and experimental randomization. The biological process of meiosis is inherently random, ensuring that multiple children from a single pair of parents have independent variation in their inherited genetics. Since the variation induced by meiosis precedes other events that may influence a particular trait and it is randomized within a trio, meiosis resembles a form of experimental randomization. This was recognized early in the development of genetics, such as in Fisher’s 1951 lecture [4]:

> “The different genotypes possible from the same mating have been beautifully randomised by the meiotic process. A more perfect control of conditions is scarcely possible, than that of different genotypes appearing in the same litter.”

In a properly randomized experiment, it is a basic result that association implies causation [2, 5, 6]. If one accepts meiosis as a valid form of experimental randomization, then genotypes associated with a phenotype can be interpreted causally by leveraging the randomization of alleles transmitted during meiosis.

Fisher’s observation that the meiotic process is a valid form of randomization is essentially correct within the single family or litter setting. However, when analyzing population-sampled trios that consist of multiple families, the analogy between a randomized experiment and meiosis does not trivially extend to the collection of families analyzed together. For instance, in a sampled population, there may be confounding between genetic variation and a trait. One reason for this is *population structure*, commonly observed in human populations [7]. In a population, the processes that gave rise to genetic variation may be related to the processes that influence a trait.

Even though a phenotype cannot, in principle, change a genotype and even though genotypes are *randomly sampled*, this does not mean that population-based studies in general contain the information needed to leverage the randomization due to meiosis. A related but distinct line of work, referred to as “Mendelian randomization” (MR), aims to identify non-genetic factors that are causal for phenotypes of interest. The challenges and underlying assumptions of MR have been variously discussed [8–10]. MR constructs instrumental variables from genotypes for population-sampled individuals but does not directly observe the meiotic process from parents to child. It is not a potential outcomes framework applied to trios where randomization is directly observed, as we consider here.

The current state of understanding is that other than in the single-family setting, therefore, association implies neither causation nor proximity to causal loci, which is why association studies are not considered to be linkage or causality studies. Genome-wide association studies (GWAS) determine statistical associations between genotypes and phenotypes without any general proof of causality. A framework has been proposed for causal inference where the parents and child have phased genomes available [11]. This framework utilizes a definition of causality based on probabilistic independence [12], which is not necessarily equivalent to the potential outcomes we employ here [3].

Here we develop a robust randomization-based causal framework for genotype-phenotype relationships by considering two *transmission-based methods* — one introduced here for arbitrarily distributed phenotypes in population-sampled trios and the other for an existing method applied to dichotomous traits in a particular type of trio study. These methods do not require phased genomes, but rather standard diploid genotypes for parents and child. We name the first approach the *transmission mean test* (TMT) and refer to the second approach under its known name, the *transmission disequilibrium test* (TDT). The TMT is a new set of methodologies developed here for both quantitative (continuous or counts) and binary phenotypes, when trios are randomly sampled from an arbitrarily structured population with other potential genotype-phenotype confounders. Such study designs occur, for example, when trios are sampled from a defined geographical region [13–15] or from a particular population [16, 17], or when trios are sampled on the basis of parent-level attributes [18]. The TDT handles trios that have an “affected” child of a binary phenotype. Previous work described the study design for the TDT as *affected-only design*, also known as *affected family-based controls*, and have developed the TDT as a statistical test of association [19–25]. We show here that the TDT is a rigorous test of causality.

The main contributions of this work are to propose the TMT and to establish that the TMT and the TDT are valid causal inference methods within the potential outcomes framework. We start by assuming that the causal and non-causal variants are probabilistically independent. We define the concept of *directly causality* and develop the TMT to detect causal genetic effects for a broad class of traits. We prove that the TDT is a causal test for dichotomous traits in the affected-only design. We then extend the causal framework of both the TMT and the TDT to the case where causal and non-causal loci may be genetically linked by developing the concept of *causal linkage*. In order to identify scenarios appropriate for these methods and how they can be used together with existing methods, we compare the proposed causal inference framework with established association methods, such as the linear mixed model for associations (LMM) [26–28], in the presence of population structure and confounding factors associated with both the trait and causal genotypes.

## 2 Theory and Methods

### 2.1 Data structure and method overview

#### Trio data

The data for an observed trio includes the child’s measured phenotype as well as the genome-wide genotypes of both parents and the child (Figure S1). The goal here is to identify the genetic loci that are causal for the measured phenotype among the children. Let *J* be the total number of trios. A sample of *J* trios will involve 3*J* distinct individuals. We assume that no individual is the parent of more than one child in the sample and no individual appears both as a parent and as a child in the sample.

Our framework starts from the unphased biallelic *single nucleotide polymorphism* (SNP) data [29], which is more common than phased data in practical settings. Let *I* be the total number of SNPs per individual. For a SNP, let *a* and *b* be the two alleles that generate three possible genotypes {*aa, ab, bb*}. We numerically encode these three genotypes by {0, 1, 2}. In the *j*th trio, *j* ∈ [1 : *J*], let *G*_*ij*_ ∈ {0, 1, 2} be the child’s *i*th genotype, 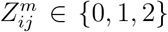 the maternal *i*th genotype, and 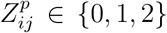 the paternal *i*th genotype, *i* ∈ [1 : *I*]. A complete set of trio genotype data includes the *I × J* matrix ***G*** = {*G*_*ij*_} for the child and *I × J* matrices 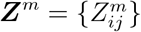 and 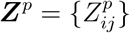 for the parents.

We additionally denote 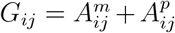 for each child, where 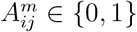 is the allele the *j*th child received from the maternal side and 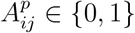 the allele from the paternal side. We do not assume that the parental transmitted alleles 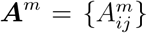 and 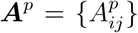 are directly observed in our framework. In addition to the genotype data, we observe a *J*-length vector of child’s phenotypes ***Y*** = {*Y*_*j*_} where *Y*_*j*_ can be either quantitative (continuous or counts) or dichotomous. Let 𝒞 be the index set of causal SNP(s) for a specific trait *Y*. The goal is to identify all *i* ∈ 𝒞 such that there exists a causal effect from *G*_*i*_ to *Y*. As 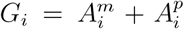 and we are not distinguishing maternal and paternal genetic effects, an equivalent aim is to estimate the causal effect from 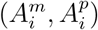 to *Y*.

#### Overview of the proposed framework

Here we give a conceptual overview of the proposed *transmission mean test* (TMT). The main idea is the characterization of the randomizations that have taken place, shown in Figure 1. Each child’s trait value is potentially subject to two independent *within-trio* randomizations, one from the maternally transmitted allele and one from the paternally transmitted allele. These transmitted alleles, 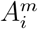 and 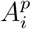, can be either 0 or 1. In order for a meaningful randomization to take place, the parent must be heterozygous. For each heterozygous parent, we assign the child to the *control* group if receiving an allele 0 and to the *treatment* group if receiving an allele 1. The control and treatment group labels are arbitrary. It is the between group difference of trait values that we need in order to estimate causal effects. The key algorithm is the assignment procedure that constructs the control and the treatment groups. If a heterozygous parent in the *j*th trio transmits an allele 0 to the child, the corresponding *Y*_*j*_ is included in the control. Conversely, if transmitting an allele 1, *Y*_*j*_ is included in the treatment. It is possible for a trait value to go into both the control and treatment groups (wherein our proposed statistic cancels out the trait value). It is also possible for a trait value to be assigned twice to the control group if both heterozygous parents contribute 0 alleles and likewise for the treatment group, as shown in Figure 1.

**Figure 1:**
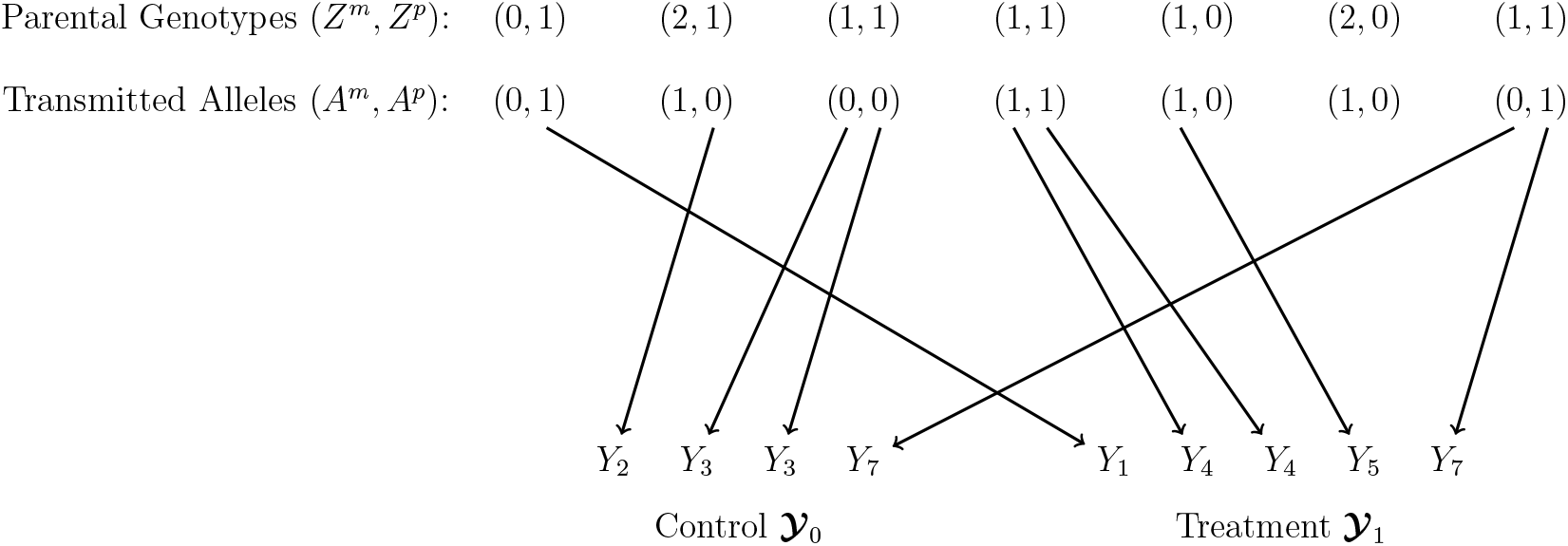
Schematic of the TMT assignment procedure. Encode the two possible transmitted alleles at the *i*th locus as allele 0 and 1. Let *Y* be the value of the child’s phenotype. In the *j*th trio, if a heterozygous parent transmits an allele 0 to the child, the corresponding phenotype *Y*_*j*_ is included in the control group. If a heterozygous parent transmits an allele 1, the corresponding *Y*_*j*_ is included in the treatment group. When there are two heterozygous parents, a trait can be included twice within one group or it can be assigned simultaneously to both groups. We do not observe a randomization from a homozygous parent.

Two vectors, **𝒴**_0_ and **𝒴**_1_, are generated, representing the observed trait values in the control and the treatment. Both **𝒴**_0_ and **𝒴**_1_ contain two types of elements: one attributed to the maternal side and the other attributed to the paternal side, written as

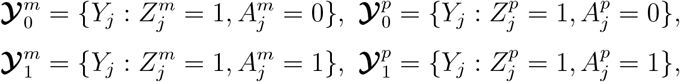

where 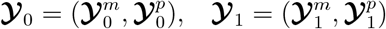.

Randomizations, when occurring in concordance with the required assumptions, allow one to straightforwardly infer causality from associations, even in the presence of confounders. This is well known in simple scenarios, such as a study with a single randomized variable that meets all required assumptions. However, the scenario we consider here with trios is non-standard and presents some interesting challenges. Sampled trios can have zero, one, or two heterozygous parents. A trio with two heterozygous parents allows us to record two randomizations and it does not leave a homozygous parent acting as a potential confounder. However, the randomization of the two parents can be in opposite directions. In the case of one heterozygous parent and one homozygous parent, it should be noted that confounders with genotype (e.g., via population structure or confounding non-genetic variation) can still enter the child’s trait value through the homozygous parent since there is no meaningful randomization from that parent. Trios with zero heterozygous parents are not used to infer causality as there are no meaningful randomizations.

We introduce a trait model and formulate it as general as possible to account for confounders. We then define a specialized potential outcomes model of causal effects, as it is nonstandard to randomly have zero, one, or two of the potential randomized variables manifesting as discernable randomizations. After these model formulations, we construct a causal effect estimand, together with a TMT statistic, based on the differences between the values in **𝒴**_0_ and **𝒴**_1_, and we prove several key operating characteristics of them.

### 2.2 Model assumptions and potential outcomes

Here, we formulate a trait model and a causal effects model for each SNP and child combination. We therefore drop the SNP subscript *i* and individual subscript *j* until they are needed.

#### The trait model

The child’s trait *Y* is modeled as:

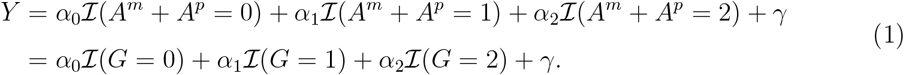

It is the case that *A*^*m*^, *A*^*p*^, *G*, and *γ* are all random variables whereas *α*_0_, *α*_1_, and *α*_2_ are fixed effects. Due to possible relatedness and population structure, the parental genotypes *Z*^*m*^ and *Z*^*p*^ may be arbitrarily dependent random variables, making *A*^*m*^ and *A*^*p*^ dependent. (We show below that some independence results on *A*^*m*^ and *A*^*p*^ hold when conditioning on certain parental genotypes *Z*^*m*^ and *Z*^*p*^.) We assume that *γ* does not interfere with the transmitted allele from heterozygote parents, detailed precisely in Assumption 2 below. This model is more general than is typically assumed for the popular linear-mixed effects approach to correct for polygenic background and population structure.

One may propose to fit the trait model in Equation (1) by a simple linear regression between the child’s trait *Y* and genotypes *G*. In order to obtain an unbiased estimated of *α*_0_, *α*_1_, and *α*_2_, such a regression approach would require *exogeneity*, i.e., 𝔼[*γ*|*G*] = 0 or equivalently ℂ(*γ, G*) = 0. However, this is not necessarily satisfied according to the assumptions underlying Equation (1). For example, when there is population structure or dependence among genetic and non-genetic effects, then exogeneity is violated.

One assumption we make about the trait model is that the genetic effects are either non-decreasing or non-increasing, but we do not assume an additive genetic model. As shown below, this allows us to tractably deal with the fact that there are two potential parental randomizations within each trio.

##### Assumption 1.

*The conditional expectation of Y given parental transmitted alleles*, 𝔼[*Y* |*A*^*m*^, *A*^*p*^], *is either a non-decreasing function of A*^*m*^ + *A*^*p*^ *such that*

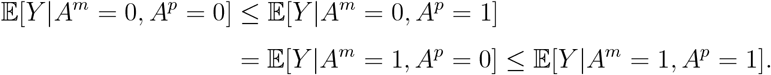

*or* 𝔼[*Y* |*A*^*m*^, *A*^*p*^] *is analogously a non-increasing function of A*^*m*^ + *A*^*p*^.

Note that under the trait model in Equation (1), if 𝔼[*Y* |*A*^*m*^, *A*^*p*^] is non-decreasing, then *α*_0_ ≤ *α*_1_ ≤ *α*_2_; if 𝔼[*Y* |*A*^*m*^, *A*^*p*^] is non-increasing, then *α*_0_ ≥ *α*_1_ ≥ *α*_2_. Since the alleles are arbitrarily coded as 0 or 1, without loss of generality we let the allele be 1 that yields an non-decrease in 𝔼[*Y* | *A*^*m*^, *A*^*p*^] and 0 otherwise. This is only for mathematical convenience and does not restrict our procedure or theory.

#### Potential outcomes

To study the causal effect of the parental transmitted alleles (*A*^*m*^, *A*^*p*^) on the child’s phenotype *Y* in standard potential outcomes parlance, we model potential outcomes as variables representing the phenotype that the child would have developed if receiving a particular allele from the parental side of interest. Let *A* be the parental transmitted allele. The potential outcomes are *Y* (*A* = 0) and *Y* (*A* = 1), sometimes simplified as *Y* (1) and *Y* (0). Then the observed trait value *Y* can be written as a function of the potential outcomes:

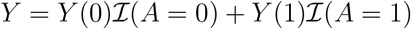

where ℐ (·) is the indicator function. Since the meiotic process for the transmission of the allele only happens once for each child, either *Y* (0) or *Y* (1) is observed, but not both. When considering a particular parental side, we write *Y* ^*m*^(0) and *Y* ^*m*^(1) for the maternal side and *Y* ^*p*^(0) and *Y* ^*p*^(1) for the paternal side. Under the trait model in Equation (1), the potential outcomes are

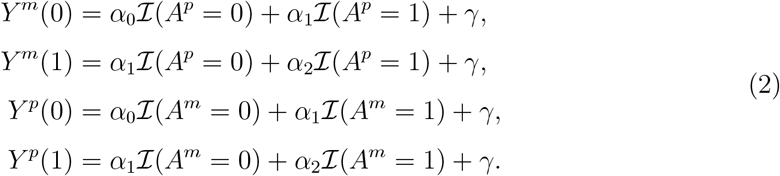

Note that these potential outcomes are on a per parent basis, while the goal is to consider both parents simultaneously. We will next define causal effects on a per parent basis and then combine them for an overall joint parent transmitted allele causal effect – or equivalently causal effect of *G* on *Y*.

### 2.3 Causal effects and regression effects

#### Average causal effect

A standard way to test for directly causality is to test for a non-zero *average causal effect* (ACE) which is a function of potential outcomes.

##### Definition 1

(ACE for an allele). *The Average Causal Effect (ACE) of a parental transmitted allele A on Y is the average difference between potential outcomes Y* (*A* = 1) *and Y* (*A* = 0): ACE(*A* → *Y*) = 𝔼[*Y* (*A* = 1)] − 𝔼[*Y* (*A* = 0)].

Under the trait model and Equation (1), notice that ACE(*A* → *Y*) = 0 if and only if *α*_0_ = *α*_1_ = *α*_2_. Otherwise, ACE(*A* → *Y*) ≠0. This allows us to define the ACE of the child’s genotype on the trait in a manner that permits us to consider both parental transmitted alleles.

##### Definition 2

(ACE for genotype). *Under the trait model and Equation* (1), *we say* ACE(*G* → *Y*) = 0 *if and only if α*_0_ = *α*_1_ = *α*_2_. *Otherwise, we say* ACE(*G* → *Y*)≠ 0.

Since *G* = *A*^*m*^ +*A*^*p*^, we will use ACE(*G* → *Y*) and ACE (*A*^*m*^, *A*^*p*^) → *Y* interchangeably. We show formally that ACE(*A* → *Y*) ≠ 0 if and only if ACE(*G* → *Y*)≠ 0 through the following lemma.

##### Lemma 1.

*Under the trait model and Assumption 1*,

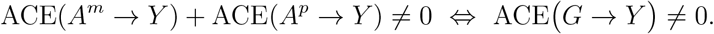

We prove Lemma 1 in Appendix A.1. By Lemma 1, a sufficient and necessary step to test *G* → *Y* is to test the sum of ACE(*A*^*m*^ → *Y*) and ACE(*A*^*p*^ → *Y*) for equality to zero or not. This allows us to use trios with only one parental heterozygote or trios with two parental heterozygotes in a single estimator and test-statistic.

##### Definition 3

(directly causality). *(A) The parental transmitted allele A is directly causal for the child’s trait Y*, *denoted A* → *Y*, *if* ACE(*A* → *Y*) ≠ 0;

*(B) The child’s genotype G is directly causal for Y*, *denoted G* → *Y*, *if* ACE *G* → *Y* ≠ 0.

#### Different between the causal effect and regression effect

Since the observed samples are drawn from the conditional distribution *Y* |*A* instead of the marginal distribution of *Y*, what we observe in reality is a conditional expectation that we address by the *average regression effect* (ARE), ARE(*Y* |*A*) = 𝔼[*Y* |*A* = 1] − 𝔼[*Y* |*A* = 0]. In general, ARE(*Y* |*A*)does not equal ACE(*A* → *Y*), which is true for the trait model in Equation (1) where the AREs are:

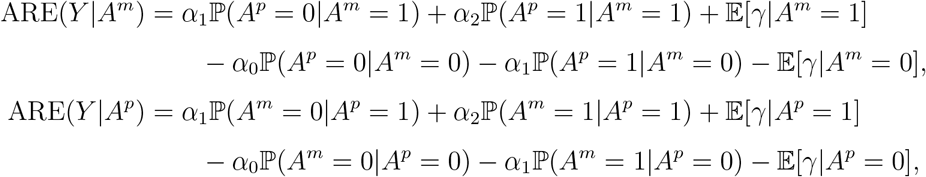

while the ACEs are:

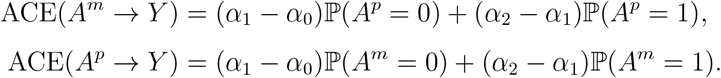

Scenarios where the ACE and ARE may be unequal include but are not limited to:

A. The dependence between *A* and *Y* is confounded by another variable such as population structure so that 𝔼[*γ*|*A* = 1] − 𝔼[*γ*|*A* = 0] ≠ 0;
B. The parental transmitted alleles *A*^*m*^ and *A*^*p*^ are not independent, which is common when population structure or relatedness exists.

Therefore, a direct regression between *Y* and *A*^*m*^ + *A*^*p*^ is usually not sufficient to detect causal effects.

### 2.4 Randomization via heterozygous parents

We develop an inference framework to estimate ACE rather than ARE. To this end, we are including the randomizations based on the parental genotypes *Z*^*m*^ and *Z*^*p*^. Under Mendel’s laws, a key fact is

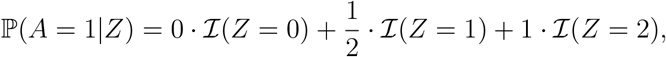

which is the probability of the child to receive allele 1 instead of allele 0 given *Z*. Here *Z* = 1 denotes a heterozygous parent who has an allele 1 and an allele 0, *Z* = 0 a homozygous parent with two alleles 0, and *Z* = 2 a homozygous parent with two alleles 1.

Notice that ℙ(*A* = 1 |*Z* = 0) = 0 and ℙ(*A* = 1 |*Z* = 2) = 1. Both cases will eliminate the possibility to observe a randomization and therefore cannot be included in estimating ACE. For the transmission from a heterozygous parent, Mendel’s Law of Segregation implies that ℙ(*A* = 1 | *Z* = 1) = 1*/*2, which indicates an equal probability to be assigned to either the control or the treatment. Therefore, both potential outcomes can be observed for trios with at least one heterozygous parent. This serves as the inclusion criteria for the TMT.

When violating Mendel’s Law of Segregation, ℙ(*A* = 1 | *Z* = 1) may be slightly shifted away from 1*/*2, which is known as “transmission distortion” [30–32]. Even though various biological processes, such as biased segregation during meiosis and differential success of gametes in achieving fertilization, may lead to a skewed value of ℙ(*A* = 1|*Z* = 1), previous research reported a shifting value for human genome *<* 0.005 genome-wide and *<* 0.0007 per locus [32]. Therefore, we assume ℙ(*A* = 1|*Z* = 1) = 1*/*2 for the current work. If there’s strong evidence for transmission distortion, one may use an estimate of ℙ(*A* = 1|*Z* = 1) to adjust the assignment probability in the TMT.

Now we state an essential assumption and lemma for estimating the ACE based on the randomization provided by alleles transmitted from heterozygous parents.

#### Assumption 2.

*Under the trait model in Equation* (1),

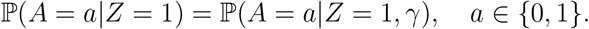

Assumption 2 follows Mendel’s Law of Segregation in that each parent transmits one allele randomly to the child during meiosis, which proceeds the development of the child’s trait.

Then both ℙ(*A* = *a*| *Z* = 1) and ℙ(*A* = *a* |*Z* = 1, *γ*) equal 1/2. We use Assumption 2 to show Lemma 2 below, which is a classical assumption of causal inference under the potential outcomes framework.

#### Lemma 2.

*Under the trait model in Equation* (1), *the potential outcomes of a child’s trait, Y* (0) *and Y* (1), *are conditionally independent of the parental transmitted allele, A, given heterozygous parental genotype, Z* = 1, *written as*

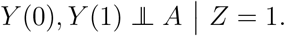

For a specific parental side, (*Y* ^*m*^(0), *Y* ^*m*^(1)) **╨** *A*^*m*^ | *Z*^*m*^ = 1 and (*Y* ^*p*^(0), *Y* ^*p*^(1)) **╨** *A*^*p*^ | *Z*^*p*^ = 1. We prove Lemma 2 in Section A.2. In this proof, we first show the independence between the maternal and paternal transmitted alleles conditional on a heterozygous parent, i.e., *A*^*p*^ **╨** *A*^*m*^|*Z*^*m*^ = 1 and *A*^*m*^ **╨** *A*^*p*^|*Z*^*p*^ = 1. This demonstrates how the randomization via heterozygous parents is independent from confounding factors such as population structure. Next we show that since the potential outcomes are functions of one parent’s transmitted alleles and *γ*, Assumption 2 can be used to complete the proof for Lemma 2.

An equivalent statement of Lemma 2 is ℙ (*A* = 1|*Y* (0), *Y* (1), *Z* = 1 = ℙ *A* = 1|*Z* = 1). Lemma 2 satisfies the unconfoundedness assumption in “intention-to-treat analysis” [3]. Intuitively, it implies that the allele transmission during meiosis is independent of the child’s trait development later. Mathematically, Lemma 2 implies: (i) the equality in distributions of (*Y* (0) | *A* = 0, *Z* = 1) and *Y* (0) *A* = 1, *Z* = 1, and (ii) the equality in distributions of (*Y* (1)|*A* = 0, *Z* = 1) and (*Y* (1)|*A* = 1, *Z* = 1). Although these two pairs of equality in distributions are not testable since we cannot observe (*Y* (0) | *A* = 1, *Z* = 1) and (*Y* (1) | *A* = 0, *Z* = 1), they are necessary for inferring causal relationships. In practice, Lemma 2 is generally satisfied if the variation induced by meiosis precedes other events that may influence a particular trait, i.e., the trait is developed after the allele transmission in meiosis.

### 2.5 TMT parameter to measure causal effects

We now define a parameter that captures the causal effect and we connect it to ACE (*G* → *Y*) through Theorem 1 shown below. Let *N* be the total number of heterozygous parents in that

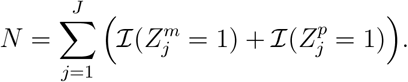

Note that *N* ≤ 2*J*. By conditioning on *N*, we derive an unbiased statistic in Section 2.6 for the following parameter.

#### Definition 4

(TMT parameter). *Let δ*_*TMT*_ *be the TMT parameter such that:*

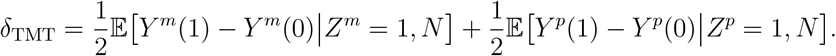

#### Theorem 1.

*Under the trait model in Equation* (1), ACE(*G* → *Y*) ≠ 0 *if and only if δ*_TMT_≠ 0.

*Proof*. The trait model implies

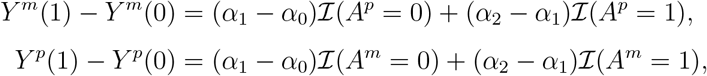

so that

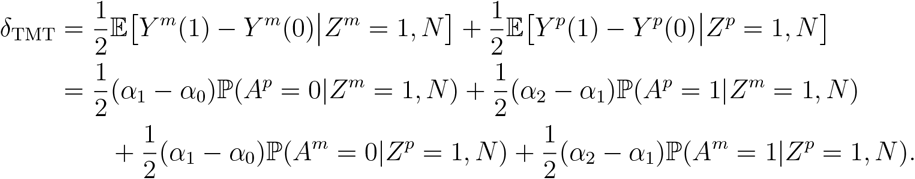

Under Assumption 1, either *α*_0_ ≤ *α*_1_ ≤ *α*_2_ or *α*_0_ ≥ *α*_1_ ≥ *α*_2_. Since 0 *<* ℙ(*A*^*p*^ = 1|*Z*^*m*^ = 1, *N*) *<* 1 and 0 *<* ℙ(*A*^*m*^ = 1|*Z*^*p*^ = 1, *N*) *<* 1, then *δ*_TMT_≠ 0 if and only if *α*_0_≠*α*_1_ or *α*_1_ ≠ *α*_2_, which is equivalent to ACE(*G* → *Y*) ≠ 0 by Definition 2.

### 2.6 Unbiased TMT estimand

We now derive an estimator of *δ*_TMT_ and show it is unbiased.

#### Assignment indices

Let *J*-length vectors ***W*** _0_ = {*W*_0*j*_} and ***W*** _1_ = {*W*_1*j*_} be assignment indices for the control and the treatment,

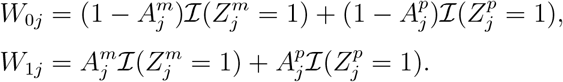

Note that *W*_0*j*_, *W*_1*j*_ ∈ {0, 1, 2}, and whenever *W*_0*j*_ = *W*_1*j*_ = 0 this implies both parents are homozygous. We summarize all possible observations and corresponding assignments in Table 1 where no ambiguous assignment exists. These assignment indices ***W*** _0_ and ***W*** _1_ are further utilized to construct the estimator of *δ*_TMT_.

**Table 1:** The TMT assignment for all possible parent-child trio observations.

| $(Z_j^m, Z_j^p, G_j)$ | $W_{0j}$ | $W_{1j}$ | $(Z_j^m, Z_j^p, G_j)$ | $W_{0j}$ | $W_{1j}$ | $(Z_j^m, Z_j^p, G_j)$ | $W_{0j}$ | $W_{1j}$ |
| --- | --- | --- | --- | --- | --- | --- | --- | --- |
| (2, 2, 2) | 0 | 0 | (1, 2, 1) | 1 | 0 | (1, 0, 0) | 1 | 0 |
| (2, 1, 2) | 0 | 1 | (1, 1, 2) | 0 | 2 | (0, 2, 1) | 0 | 0 |
| (2, 1, 1) | 1 | 0 | (1, 1, 1) | 1 | 1 | (0, 1, 1) | 0 | 1 |
| (2, 0, 1) | 0 | 0 | (1, 1, 0) | 2 | 0 | (0, 1, 0) | 1 | 0 |
| (1, 2, 2) | 0 | 1 | (1, 0, 1) | 0 | 1 | (0, 0, 0) | 0 | 0 |

#### TMT estimand

To estimate *δ*_TMT_, we start from a preliminary statistic and prove its unbiasedness as follows.

##### Definition 5

(Preliminary statistic). *Define the preliminary statistic* 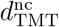 *as the sample mean difference between the treatment and the control, i*.*e*.,

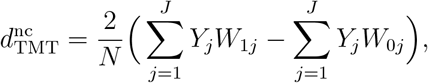

*where the superscript “nc” stands for “not-centered”*.

##### Lemma 3.

*The statistic* 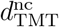 *is unbiased given N in that*

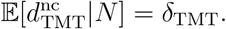

We prove Lemma 3 in Section A.3. However, 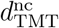 includes unnecessary variation in that the means of *Y*_*j*_ are perturbed by the random variables *W*_0*j*_ and *W*_1*j*_. One may then center each *Y*_*j*_ by its overall mean and reduce the variance of 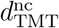 by forming the centered statistic.

##### Definition 6

(Centered statistic). *Let µ*_0_, *µ*_1_ *be the population means of potential outcomes Y* (0), *Y* (1) *conditioning on heterozygous parents such that:*

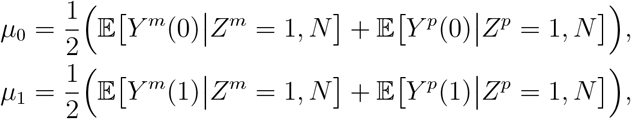

*and their average value*

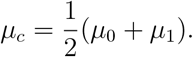

*Define the centered statistic as*

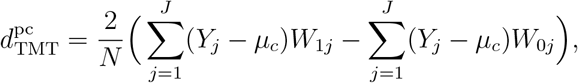

*where the superscript “pc” stands for “population-centered”*.

When *µ*_*c*_ is unknown, we form an unbiased estimate 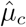, through which we further form the ultimate TMT statistic.

##### Definition 7

(TMT statistic). *Define the population mean estimate* 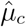 *such that*

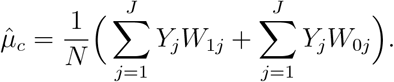

*The ultimate TMT statistic used in practice involving no unobserved parameters, which we will denote by d*_TMT_, *is*

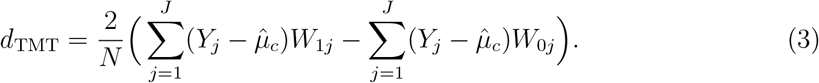

##### Lemma 4.

*The statistic* 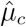 *is unbiased for µ*_*c*_ *in that*

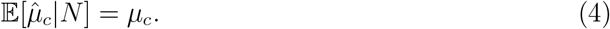

*Also*,

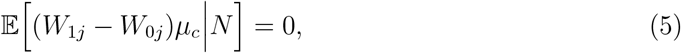

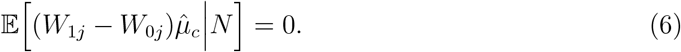

We prove Equation (4) in Section A.4. Equation (5) and Equation (6) are proved in Section A.5 and are used to show that both centered statistics are unbiased.

##### Lemma 5.

*Under the trait model in Equation* (1), *the statistics* 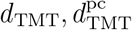 *are unbiased for δ*_TMT_ *in that*

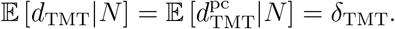

We prove Lemma 5 in Section A.6. Calculating *d*_TMT_ based on observed trio data enables the assessment of the existence of causality between the locus of interest and the target trait, which will be discussed in Section 2.8. Before that, we will need an estimate of the variance of *d*_TMT_ in order to construct a test-statistic.

### 2.7 Sampling variance of the TMT estimand

Deriving a sampling variance estimate and proving its operating characteristics is challenging in this setting. It has some properties that are different from a traditional randomized study. Note that when there is population structure or relatedness, the parental genotypes 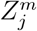 and 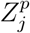 may be dependent for each family *j*. Also, *Z*_*j*_ and *Z*_*k*_ may be dependent for any two families *j*≠ *k* and for any combination of maternal and paternal genotypes. This means that unconditionally, the potential outcomes 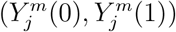 and 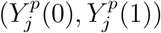 may be dependent within each family *j*. This follows because these two pairs of potential outcomes are functions of the parental transmitted alleles 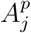 and 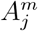, respectively; they are therefore functions of the parental genotypes 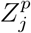 and 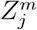, respectively. Likewise, (*Y*_*j*_(0), *Y*_*j*_(1)) and (*Y*_*k*_(0), *Y*_*k*_(1)) may be dependent between families *j*≠ *k* for any combination of maternal and paternal genotypes.

We showed that 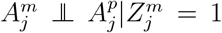 and 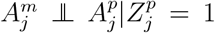 via Equation (A2.c) from the proof of Lemma 2. Thus, within each family *j*, the potential outcomes are independent conditional on a heterozygous parent. In deriving the sampling variance here, we show several results that provide sufficient properties regarding between family covariances.

Another property of our setting that is different from a traditional randomized study is that the number of individuals assigned to treatment or control is random, whereas it would be predetermined in a traditional setting [3]. Further, a child can be assigned twice to treatment or control. Therefore, when deriving our sampling variance estimate, we first condition on these random events and then extend the estimate to take into account our particular randomization.

#### Sampling variance estimate

To develop a test based on *d*_TMT_, we first form a sampling variance estimate of the test-statistic. Let 𝒥 = {1, 2, …, *J*}. Define the following sets:

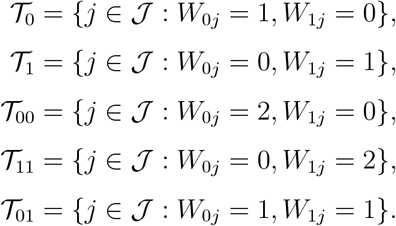

Notice that |𝒯_0_| + |𝒯_1_| + 2|𝒯_00_| + 2|𝒯_11_| + 2|𝒯_01_| = *N*. Note that for *j* ∈ 𝒯_01_, (*W*_1*j*_ −*W*_0*j*_)*Y*_*j*_ = 0 so these observations do not appear in *d*_TMT_. Denote means and variances of the other four sets as follows:

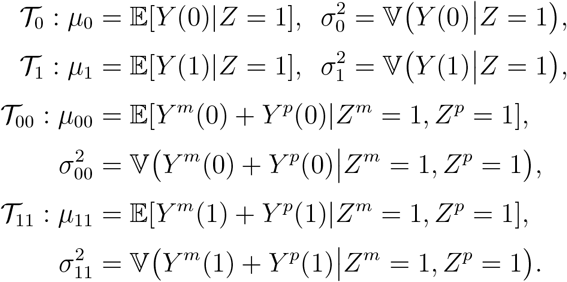

##### Definition 8

(Sampling variance estimate). *We form the following estimates utilized in the ultimate sampling variance estimate*.

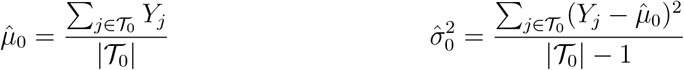

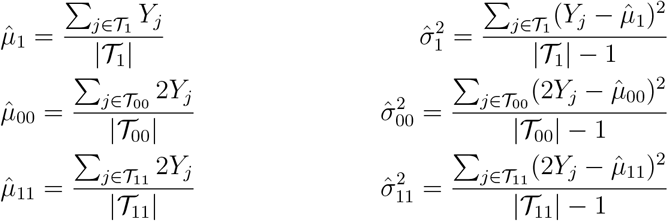

*From these, we construct the sampling variance estimate for d*_TMT_ *as:*

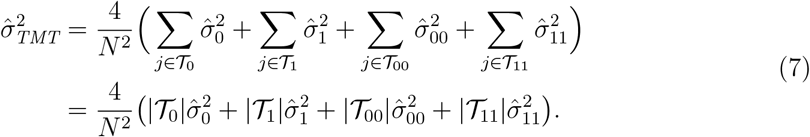

#### Unbiasedness of the sampling variance estimate

We first discuss the expected value 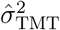 as an estimate for 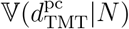 by Lemma 6 to Lemma 9. Then we show 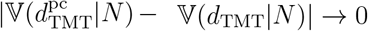 with probability 1 as *J* → ∞ in Lemma 10.

##### Lemma 6.

*The estimate* 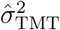 *is unbiased for*

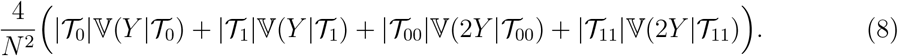

Lemma 6 follows the fact that the sampling variance estimands 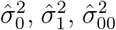, and 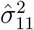 are unbiased estimators for their corresponding population variances 𝕍(*Y* |𝒯_0_), 𝕍(*Y* |𝒯_1_), 𝕍(2*Y* |𝒯_00_), and 𝕍(2*Y* |𝒯_11_) [3].

We now show that Equation (8) is the variance of 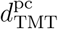 when *δ*_TMT_ = 0 and it is a lower bound when *δ*_TMT_ ≠0. To do so, we detail an assumption as follows.

##### Assumption 3.

*Consider a sample of* 2*J parents including N heterozygous parents*.

A. *N/*(2*J*) → ℙ(*Z* = 1) *with probability 1 when J* → ∞;
B. *For j, k* ∈ 𝒥 *and j*≠ *k*, ℙ(*Z*_*j*_ = *a, Z*_*k*_ = *b*|*N*) → ℙ(*Z*_*j*_ = *a, Z*_*k*_ = *b*) *with probability 1 when J* → ∞ *for a, b* ∈ {0, 1, 2};
C. *For j, k* ∈ 𝒥 *and j*≠ *k*, ℂ(*Z*_*j*_, *Z*_*k*_) ≥ 0.

Note that Assumption 3(B) and Assumption 3(C) hold for any combination of maternal or paternal genotypes. By the Law of Total Probability, Assumption 3(B) implies that ℙ(*Z* = *a*|*N*) → ℙ(*Z* = *a*) with probability 1 when *J* → ∞ for *a* ∈ {0, 1, 2}. Also note that Assumption 3(C) is satisfied in prevalent population genetics models including the Identical-By-Descent (IBD) model, the co-ancestry model, etc [33].

##### Lemma 7.

*Under Assumption 3*, *for j, k* ∈ 𝒥 *and j* ≠ *k*,

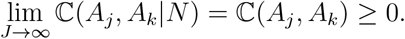

*This holds for any combination of maternal or paternal genotypes*.

We prove Lemma 7 in Section A.7 and use it to prove the following lemma.

##### Lemma 8.

*Write D*_*j*_ = (*W*_1*j*_ − *W*_0*j*_)(*Y*_*j*_ − *µ*_*c*_). *The covariance between families*

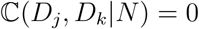

*for j, k* ∈ 𝒥 *and j k and when δ*_TMT_ = 0. *Let ω* = ℙ(*Z* = 1), *which is the proportion of heterozygous parents. When δ*_TMT_ ≠ 0, *as J* → ∞, *the above covariance converges to a non-negative value in that*

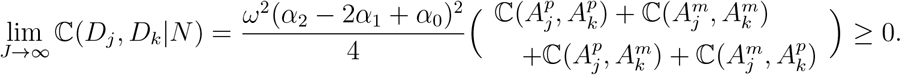

Note that *α*_0_, *α*_1_ and *α*_2_ are parameters for the trait model in Equation (1). We prove Lemma 8 in Section A.8 and use it to derive the following lemma.

##### Lemma 9.

*Under the trait model in Equation* (1) *with arbitrary relatedness between families in a trio study:*

A. *When* 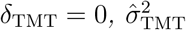 *is an unbiased estimator for* 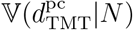.
B. *When* 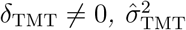 *underestimates* 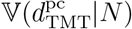 *as J* → ∞.

We prove Lemma 9 in Section A.9. Lemma 9 justifies 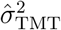 as an unbiased estimator for the variance of 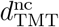 when *δ*_TMT_ = 0 and a lower bound for the variance of 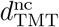 when *δ*_TMT_≠ 0. These properties will allow us to derive a valid hypothesis test of *δ*_TMT_ = 0 versus *δ*_TMT_ 0 below. To further connect these results back to *d*_TMT_, we derive the following lemma.

##### Lemma 10.

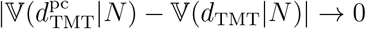 *with probability 1 as J* → ∞.

We prove Lemma 10 in Section A.10. This implies that, as 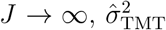 is consistent for the variance of *d*_TMT_ when *δ*_TMT_ = 0 and is a lower bound for the variance of *d*_TMT_ when *δ*_TMT_≠ 0.

### 2.8 Proposed hypothesis test

Here we derive a hypothesis test where the null hypothesis is *H*_0_ : *δ*_TMT_ = 0 and the alternative hypothesis is *H*_1_ : *δ*_TMT_ ≠ 0. Recall Theorem 1, where it was shown that *δ*_TMT_ ≠ 0 if and only if ACE(*G* → *Y*) ≠ 0, implying the hypothesis test is the desired test of causality. Under the trait model in Equation (1), the null hypothesis of no causality is true if and only if *α*_0_ = *α*_1_ = *α*_2_. Since *d*_TMT_ is unbiased for *δ*_TMT_ in Section 2.6, we form a test-statistic based on *d*_TMT_. The square root of the sampling variance estimate 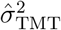 developed in Section 2.7 is also utilized as the standard error, which allows us to form a test-statistic with a known null distribution.

We propose the following test-statistic:

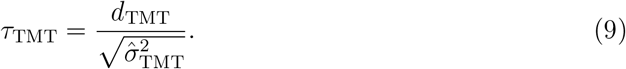

By the Central Limit Theorem, *τ*_TMT_ is approximately Normal(0, 1) when the null hypothesis, *H*_0_ : *δ*_TMT_ = 0, is true. The p-value is calculated by

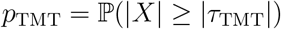

where *X* ∼ Normal(0, 1). Rather than using the Normal(0, 1) distribution to calculate a

*p*-value, one could also use a permutation null distribution. Details for this permutation test are in Section B.3. We will show later that the permutation null and the Normal(0, 1) are similar in practice (Section 3.1).

### 2.9 TDT is a test of causality

The TDT is applied when trios are recruited by identifying children with a particular disease. This sampling strategy is natural when studying a disease commonly contracted in childhood [34–36]. As it is required to diagnose in advance whether a child has the disease or not, the TDT handles binary traits. Children who are sampled for TDT analysis have trait value *Y*_*j*_ = 1. Denoting the number of transmitted allele 0 by *N*_0_ and allele 1 by *N*_1_,

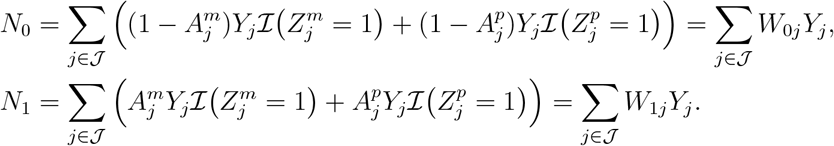

The above procedure is visualized in Figure 2. One then tests for a significant difference between *N*_1_ and *N*_0_. In the standard implementation [21], McNemar’s test [37] is performed based on a *χ*^2^ statistic

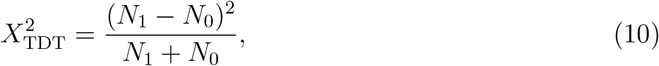

with 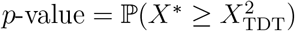 where *X*^*^ has a 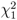 distribution.

**Figure 2:**
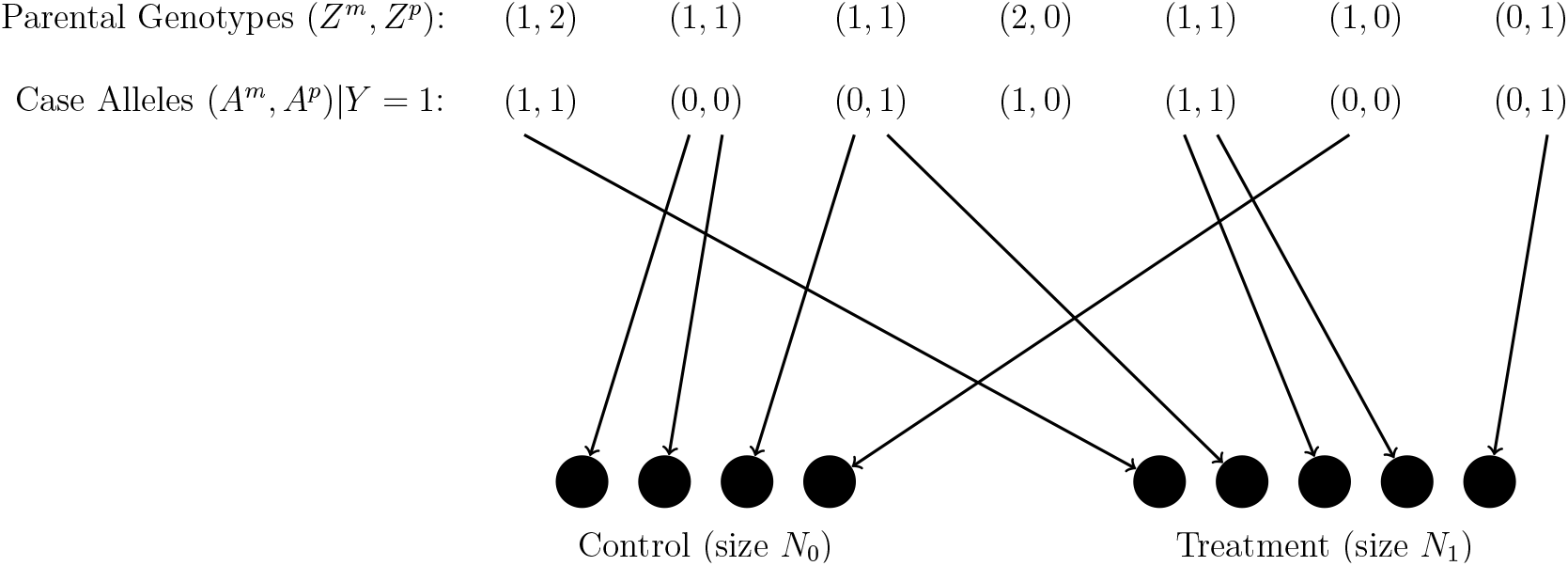
Schematic of the TDT assignment procedure. If a heterozygous parent transmits an allele 0 to the affected child, an event (denoted by a solid dot) is recorded in the control. Alternatively, if transmitting an allele 1, an event is recorded in the treatment.

We show that the TDT is a test of causality via the following theorem, which relies on our causal inference framework for the TMT. We then show below that a component of the TDT statistic is an unbiased estimator of the causal effect analogous to *d*_TMT_.

#### Theorem 2.

*The null hypothesis of TDT is true if and only if* ACE *G* → *Y* = 0.

*Proof*. We first construct a parameter analogous to the TMT parameter *δ*_TMT_ under the TDT setting by defining a TDT parameter *δ*_TDT_, which involves a factor that accounts for the affected-only sampling. Suppose the affected children are sampled from a population where the disease prevalence given parental heterozygote is

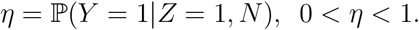

Note that *η* is not needed for the test but just for bridging the TDT and the TMT.

#### Definition 9

(TDT parameter). *Define the TDT parameter δ*_TDT_ *such that*

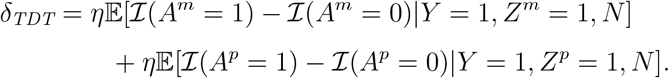

A relevant difference between the TDT and the TMT for dichotomous traits is that the original TDT only samples trios with the child’s trait value *Y*_*j*_ = 1, whereas the causal framework for the TMT is based on trios with the child’s trait value *Y*_*j*_ = 1 and *Y*_*j*_ = 0 randomly sampled from the underlying population. To fit the TDT under the causal framework for the TMT, we derive causal properties of the TDT with respect to the underlying population with both *Y*_*j*_ = 1 and *Y*_*j*_ = 0. We explain the connection between *δ*_TDT_ and *δ*_TMT_ as the following.

#### Lemma 11.

*When the trait only takes two values, such that Y* ∈ {0, 1}, *then δ*_*TDT*_ = *δ*_*TMT*_.

We prove Lemma 11 in Section A.11. It implies the equivalence between *δ*_TDT_ and *δ*_TMT_ for binary traits. It enables developing an estimand for *δ*_TDT_ in a similar format as the TMT estimand *d*_TMT_. To this end, we consider the following unbiased TDT estimand.

#### Definition 10

(TDT estimand). *The TDT estimand is defined as:*

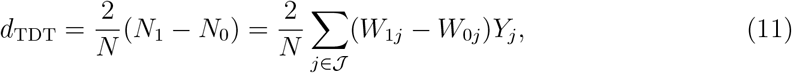

*where Y*_*j*_ = 1 *for all j* ∈ 𝒥.

Note that *d*_TDT_ = *d*_TMT_ when *Y*_*j*_ = 1 for all *j* ∈ 𝒥.

#### Lemma 12.

*Under the trait model in Equation* (1), *d*_TDT_ *is unbiased for δ*_TDT_ *in that*

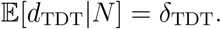

We prove Lemma 12 in Section A.12.

To connect *d*_TDT_ to the original TDT test-statistic, consider 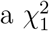, which is the difference between the proportion of alleles 1 (*N*_1_*/N*) and the proportion of alleles 0 (*N*_0_*/N*) transmitted from affected child’s heterozygous parents. The theory of McNemar’s statistic [37] utilizes the following component

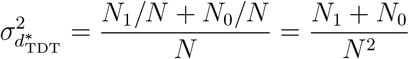

to derive a test-statistic that equals *χ*^2^ with one degree of freedom as

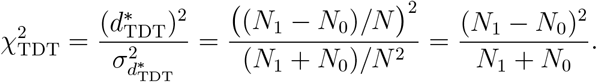

Therefore, the original TDT is equivalent to testing *H*_0_ : *δ*_TDT_ = 0 vs. *H*_1_ : *δ*_TDT_ ≠ 0. By Lemma 11, *δ*_TDT_ ≠ 0 if and only if *δ*_TMT_ ≠ 0. By Theorem 1, *δ*_TMT_ ≠ 0 if and only if ACE *G* → *Y* ≠ 0. Thus *δ*_TDT_ ≠ 0 if and only if ACE(*G* → *Y*) ≠ 0, which completes the proof for Theorem 2.

### 2.10 Detecting causal linkage via the TMT and TDT

Here, we consider the case where a genetic marker is linked to a causal variant, due to population-level linkage disequilibrium, within-trio meiotic genetic linkage, or both. Suppose that marker *c* is causal while non-causal marker *d* is linked to *c*. Since *c* is causal the trait model from Equation (1) is *Y* = *α*_*c*0_ℐ(*G*_*c*_ = 0) + *α*_*c*1_ℐ(*G*_*c*_ = 1) + *α*_*c*2_ℐ(*G*_*c*_ = 2) + *γ*_*c*_, where the assumptions required for the TMT and TDT hold. A key assumption is Assumption 2, where it is assumed that ℙ(*A*_*c*_ = *a*|*Z*_*c*_ = 1) = ℙ(*A*_*c*_ = *a*|*Z*_*c*_ = 1, *γ*_*c*_) for *a* ∈ {0, 1}. For a causal locus, this is a plausible assumption as explained when that assumption was introduced. However, for locus *d* where *G*_*c*_ and *G*_*d*_ may be dependent, the assumption when modeling *Y* in terms of *G*_*d*_ may be violated.

We can write the trait model in terms of both genotypes *G*_*c*_ and *G*_*d*_ as

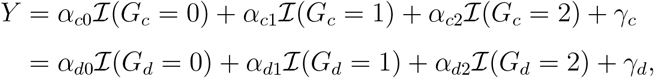

where

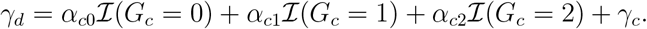

Since *c* is causal it follows that *α*_*c*1_ −*α*_*c*0_ ≠ 0 or *α*_*c*2_ −*α*_*c*1_ ≠ 0. Since *d* is not causal, it should be the case that *α*_*d*0_ = *α*_*d*1_ = *α*_*d*2_ = 0. In order to investigate the behavior of the TMT and TDT applied to marker *d*, we need to determine if ℙ(*A*_*d*_ = *a*|*Z*_*d*_ = 1) = ℙ(*A*_*d*_ = *a*|*Z*_*d*_ = 1, *γ*_*d*_) for *a* ∈ {0, 1}. (Note that when *Z*_*d*_ ≠ 1, then the transmission from this parent is not included in the TMT or TDT.) This may not be the case since *G*_*c*_ and *G*_*d*_ are dependent, so *A*_*c*_ and *A*_*d*_ may be dependent. Further, *γ*_*d*_ is a function of *A*_*c*_ (and *G*_*c*_).

For a given parent, the dependence between *A*_*d*_ and *γ*_*d*_ is driven by *A*_*c*_, so we will determine if ℙ(*A*_*d*_ = *a*|*Z*_*d*_ = 1) = ℙ(*A*_*d*_ = *a*|*Z*_*d*_ = 1, *A*_*c*_) for *a* ∈ {0, 1} by specifically calculating ℙ(*A*_*d*_ = *a, A*_*c*_ = *b*|*Z*_*d*_ = 1) for *a, b* ∈ {0, 1}. When *Z*_*c*_ = 0, then *A*_*c*_ = 0 with probability 1 and when *Z*_*c*_ = 2, then *A*_*c*_ = 1 with probability 1. Thus,

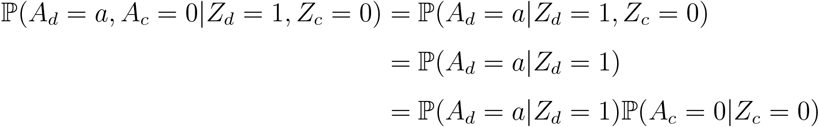

for *a* ∈ {0, 1}. The analogous result holds when *Z*_*c*_ = 2. One can conclude from this that when *Z*_*c*_ = 0 or *Z*_*c*_ = 2, then the required assumption to validly test marker *d* for causality holds for this parent-child combination.

This leaves the case of *Z*_*c*_ = 1. A given parent has two haplotypes from among the four possible haplotypes, {(0, 0), (0, 1), (1, 0), (1, 1)}. Let *H*_*cd*_ be the set of two haplotypes of the parent. Given that *Z*_*c*_ = 1, *Z*_*d*_ = 1, the possible haplotypes are *H*_*cd*_ = {(0, 0), (1, 1)} or *H*_*cd*_ = {(0, 1), (1, 0)}. Let

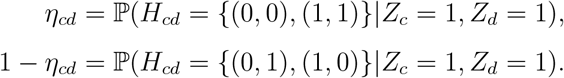

Also, let *θ*_*cd*_ be the meiotic recombination rate between loci *c* and *d*. It then follows that

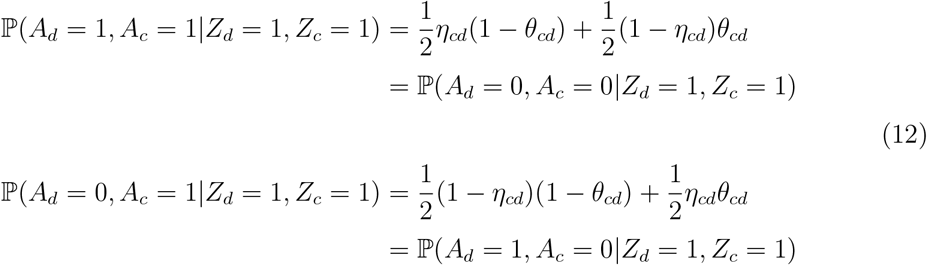

Based on Equation (12), it is trivial to show that ℙ(*A*_*c*_ = *a*|*Z*_*c*_ = 1, *Z*_*d*_ = 1) = 1*/*2 for *a* ∈ {0, 1} regardless of the values of *η*_*cd*_ and *θ*_*cb*_. We have also shown earlier that ℙ(*A* = *a*|*Z* = 1) = 1*/*2 for *a* ∈ {0, 1} at any locus. It is therefore the case that

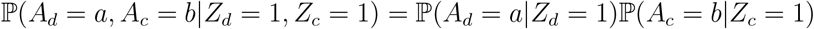

for *a, b* ∈ {0, 1} whenever the quantities in Equation (12) equal 1*/*4, which holds if and only if either *η*_*cd*_ = 1*/*2 or *θ*_*ab*_ = 1*/*2. Note that the it does not need to be the case that *η*_*cd*_ = *θ*_*ab*_ = 1*/*2; only one of the parameters needs to equal 1*/*2.

When markers *c* and *d* are in linkage equilibrium, it will be the case that *η*_*cd*_ = 1*/*2, and when markers *c* and *d* are genetically unlinked (in the meiotic sense), it will be the case that *θ*_*ab*_ = 1*/*2. When *both* of these do not hold, then the randomization at marker *d* is probabilistically dependent with the randomization at *c* and we say that these markers have *randomization linkage*. This implies that the null hypothesis of marker *d* is not true when marker *c* is causal. In this case we say that *d* is in *causal linkage* with *Y*, formalized in the following definition.

#### Definition 11

(Causal linkage of alleles). *Suppose marker c is directly causal for Y*, *where A*_*c*_ *satisfies Definition 3 in that* ACE(*A*_*c*_ → *Y*)≠ 0. *Locus d is in “causal linkage” with the trait Y if it has both linkage disequilibrium and meiotic genetic linkage with marker c. We denote causal linkage by A*_*d*_ ↪ *Y*.

If an allele *A* is itself directly causal for *Y*, then this allele *A* is trivially in causal linkage with *Y* since it is in complete linkage with itself. Thus, *A* ↪ *Y* ⇒ *A* ↪ *Y*.

#### Definition 12

(Causal linkage of genotypes). *For the child’s genotype* 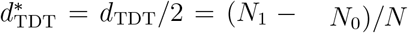, *we say* 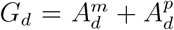 *if either* 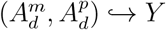 *or* 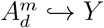.

#### Lemma 13.

*For the TMT, δ*_TMT_ ≠ 0 *at marker d only if* 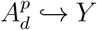. *For the TDT, δ*_TDT_≠ 0 *at marker d only if* 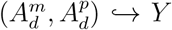.

We prove Lemma 13 in Section A.13. It enables the identification of causal linkage by testing non-zero *δ*_TMT_ via the TMT and non-zero *δ*_TDT_ via the TDT. The test-statistics *d*_TMT_ and *d*_TDT_ can be utilized to characterize the signal of causal linkage by genome-wide TMT or TDT profiles shown in Section 3.3. Under simulations reflecting levels of linkage disequilibrium observed in humans, it does not appear that causal linkage is an impediment to identifying causal loci. Under the theoretical calculations done above, it can be seen that the power is greater at *c* relative to *d*, and this is what we also observe empirically. It has been suggested that under Haldane’s model of recombination one can break the genome into independent blocks, which would allow us to test for causality at the resolution of these blocks [11]. However, empirical evidence suggests that Haldane’s model of recombination does not hold in the human genome [38–40].

## 3 Numerical Results

### 3.1 Evaluation of the TMT as a test of causality

#### Simulating trio genotypes

We simulated trio data to evaluate the accuracy and power of the TMT. We sampled the parental genotypes ***Z*** from a structured population based on a standard admixture model [33, 41, 42] of four intermediate sub-populations (Section B.1). We followed an established pipeline [33] to simulate intermediate subpopulations while keeping the overall *F*_ST_ = 0.2. (See Figure S2 for a visualization of pairwise relatedness in the structured population through co-ancestry coefficients.) We then simulated the alleles 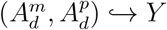 by randomly drawing one transmitted allele from corresponding parental genotypes 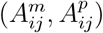, yielding the child’s genotype 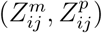.

#### Simulating child’s phenotypes

To simulate the child’s trait **𝒴** = {*Y*_*j*_} (see the schematic in Figure S3), we implemented a trait model with genetic effects from multiple loci in Equation (13). Let 𝒞 be the index set of causal loci. The traits are generated according to:

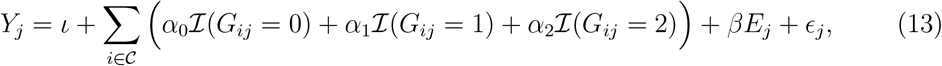

where (*ι, β*) are scalar values, (*α*_0_, *α*_1_, *α*_2_) are genetic effects per locus (made equal for all causal loci), and *ϵ*_*j*_ is the independent non-genetic random variation drawn from 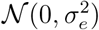. The random variable *E*_*j*_ is the non-genetic variation constructed to also be a function of the population structure. Therefore, *E*_*j*_ is a confounder between the genotypes and the trait, and the parameter *β* allows us to modulate the effect size of this confounder. Simulation details about *E*_*j*_ are summarized in Section B.2, where we also show that Equation (13) satisfies the trait model in Equation (1). Note that the ratio (*α*_2_ − *α*_1_)*/*(*α*_1_ − *α*_0_) can be any numerical value under the trait model in Equation (1). When (*α*_2_ − *α*_1_)*/*(*α*_1_ − *α*_0_) = 1, it is equivalent to an additive polygenic trait model. We display the result for (*α*_2_ − *α*_1_)*/*(*α*_1_ − *α*_0_) = 1 in Figure 3 and other (*α*_2_ − *α*_1_)*/*(*α*_1_ − *α*_0_) values in Figure S4. We set *ι* = 100 and note that any value of *ι* should work because *ι* would be subtracted from *Y*_*j*_ after centering by 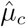 in Equation (3).

**Figure 3:**
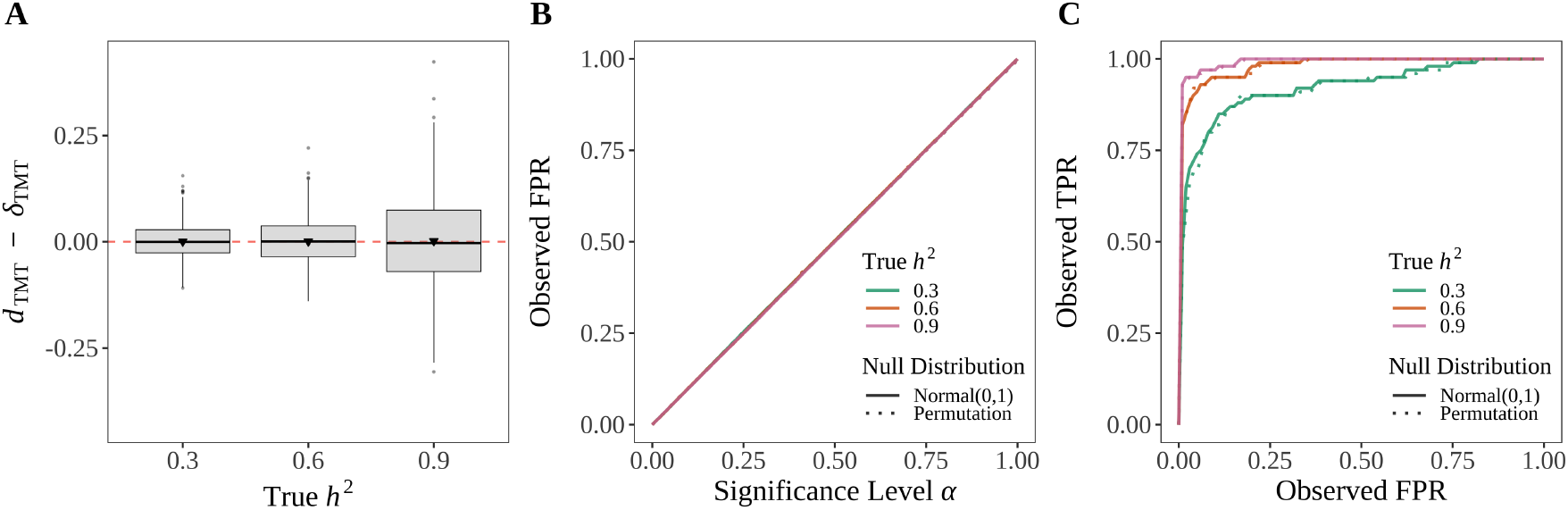
The TMT is a valid test of causality. **(A)** *d*_TMT_ is an unbiased estimator of *δ*_TMT_. We followed Section B.1 to generate 5,000 trios from a structured population (*F*_*ST*_ = 0.2) with 100,000 SNPs per individual and we randomly chose 100 loci to be causal. We simulated the child traits at the heritability levels (*h*^2^ ∈ {0.3, 0.6, 0.9}) based on Section B.2. Shown are 1000 randomly chose instances of *d*_TMT_ − *δ*_TMT_ for each heritability level. **(B)** The TMT controls the FPR at the desired significance level per experiment. **(C)** The ROC curve from a randomly chosen simulated data set. Both the Normal(0, 1) (solid lines) and permutation (dotted lines) null distributions are shown.

#### Point estimation of ACE

We numerically confirmed the accuracy of *d*_TMT_ as an unbiased estimate of the TMT parameter *δ*_TMT_ to detect non-zero ACE in Definition 1 across various values of *h*^2^ (Figure 3A). The unbiasedness of *d*_TMT_ is immune to the confounding effects from the population structure and non-genetic factors that are associated with the population structure.

#### Causal FPR and the ROC curve

We performed the TMT at each locus and calculated *p*-values for both causal and non-causal SNPs by utilizing both the Normal(0, 1) theoretical null distribution and the permutation null. We then computed the false positive rate (FPR) and the true positive rate (TPR) for causal loci at various significance thresholds between 0 and 1. The TMT strictly controls the FPR at the desired significance level across all levels of *h*^2^ (Figure 3B). The receiver operating characteristic (ROC) curve demonstrates the TMT having statistical power for detecting directly causal effects for all levels of *h*^2^, especially when *h*^2^ is high (Figure 3C). Besides the standard Normal(0, 1) null distribution, we also used a permutation null distribution to calculate a *p*-value by following Section B.3. The two null distributions show similar performance (Figure 3B and C).

#### Validation of theoretical null distribution

We started from a small sample of 100 trios and permuted **𝒴** to generate the null distributions of the test-statistic *τ*_TMT_ from Equation (9). Our results validated that *τ*_TMT_ follows the Normal(0, 1) distribution (Figure S5). Considering that population-based trio studies usually contain at least a few hundred families, the Central Limit Theorem underlying the Normal(0, 1) null distribution for *τ*_TMT_ seems to be applicable.

### 3.2 Evaluation of the TDT as a test of causality

As with the TMT evaluation, we followed the trio genotypes simulation pipeline in Section B.1 to simulate 10,000 trios with 100,000 SNPs per individual. Among the child geno-types ***G*** = {*G*_*ij*_}, we randomly chose 100 directly causal loci. To simulate a dichotomous trait, we first generated a continuous latent variable ***L*** = {*L*_*j*_} by

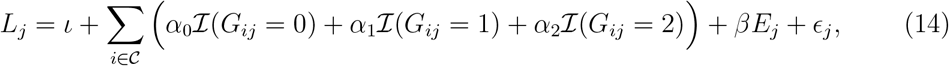

where the parameter configuration is the same as Equation (13) used for the TMT evaluation. We then used the probit model (see Section B.4) to convert *L*_*j*_ into a dichotomous trait *Y*_*j*_ according to a chosen disease prevalence. Children with *Y*_*j*_ = 0 are labeled *normal* and those with *Y*_*j*_ = 1 are labeled *affected*.

We applied the TDT on a per locus basis for trios with affected children and we also applied the TMT to a random subset of trios to match the sample size with the TDT. We computed the FPR and TPR across the range of significance levels. The TDT statistic *d*_TDT_ also demonstrates it is an unbiased estimate of the ACE for binary traits (Figure 4A). At all significance levels, the TDT controls the FPR at the desired level (as does the TMT) across all values of *h*^2^ (Figure 4B). The ROC curves demonstrate similar and reliable statistical power of the TDT and the TMT for idenitifying causal loci (Figure 4C).

**Figure 4:**
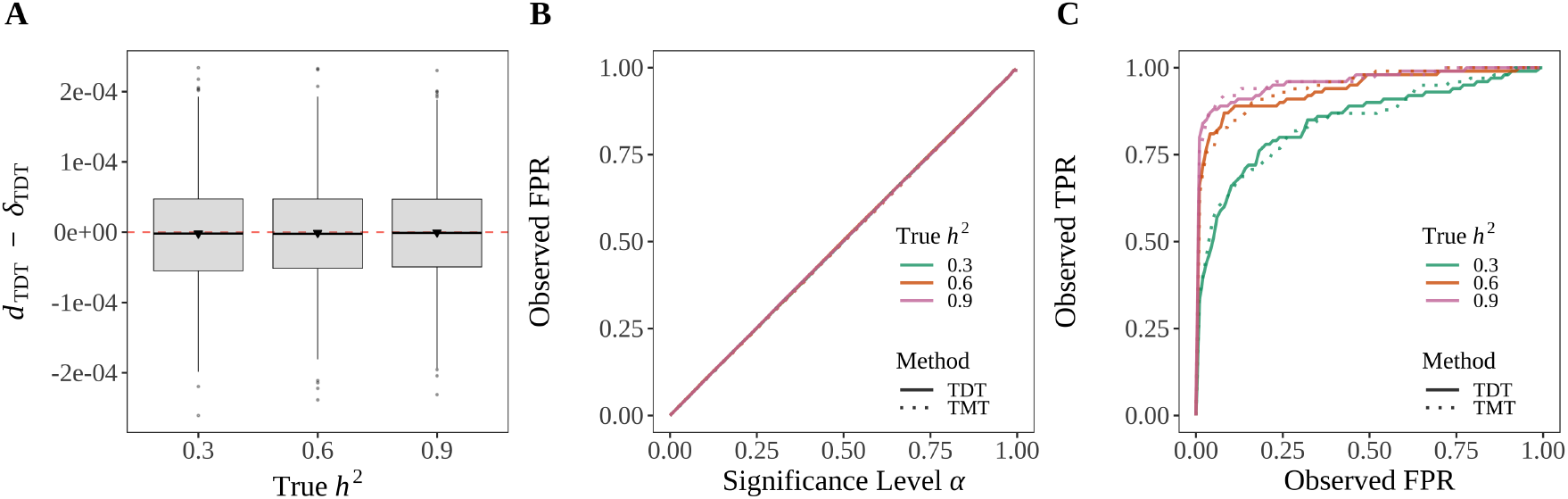
The TDT is a valid test of causality. **(A)** *d*_TDT_ is an unbiased estimator of *δ*_TDT_. We simulated 5000 affected-only trios based on the description in the text. We then calculated *d*_TDT_ and performed the TDT. Each box plot shows 1000 randomly chosen instances of *d*_TDT_ − *δ*_TDT_ at each heritability level. **(B)** The TDT and TMT control FPR at the desired significance level. We additionally applied the TMT to 5000 randomly chosen trios composed of an equal number of normal and affected children. We calculated the FPR for the TDT and TMT across a range of thresholds as shown. **(C)** The ROC curve for a randomly chosen data set for both the TDT and TMT.

### 3.3 Genome-wide TMT profile and causal linkage

To empirically validate the TMT in the presence of causal linkage, we used the software msprime [43] to simulate trio genotypes with linkage between SNPs. We generated a sample of 5,000 trios with 100,000 SNPs across 22 chromosomes per individual based on the American Admixture model [44] (simulation details in Section B.5). The level of linkage-disequilibrium (LD) in our simulated genotypes (Figure S6) matches previous findings observed in the human genome [45, 46].

We first simulated a quantitative trait for the child phenotype to implement the TMT. Among the child genotypes ***G***, we randomly chose 10 causal SNPs denoted by set *C*. We generated the child phenotypes ***Y*** by the polygenic trait model 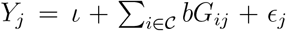 where we drew *ϵ*_*j*_ from Normal 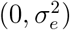 and we set 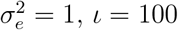. The coefficient for causal SNPs, *b*, is determined such that 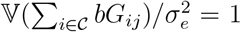 (i.e., the heritability *h*^2^ = 0.5). We displayed the TMT statistic *d*_TMT_ and corresponding *p*-values across all 100,000 SNPs as the genome-wide TMT profile (Figure 5A). To compare with linkage-equilibrium (LE), we permuted trio genotypes to remove genetic linkage, regenerated the child trait using the same parameters in the polygenic trait model above, and conducted the TMT on a per locus basis to derive *d*_TMT_ and *p*-values (Figure 5B). Under both LD and LE scenarios, the TMT shows accurate ACE estimation at the causal loci and yields a Uniform(0, 1) distribution of the *p*-values when the null hypothesis of non-causality is true. We also simulated a dichotomous trait and calculated the TDT statistic *d*_TDT_ and corresponding *p*-values to construct the genome-wide TDT profile under both LD and LE in Figure S7, where equivalent results were observed.

**Figure 5:**
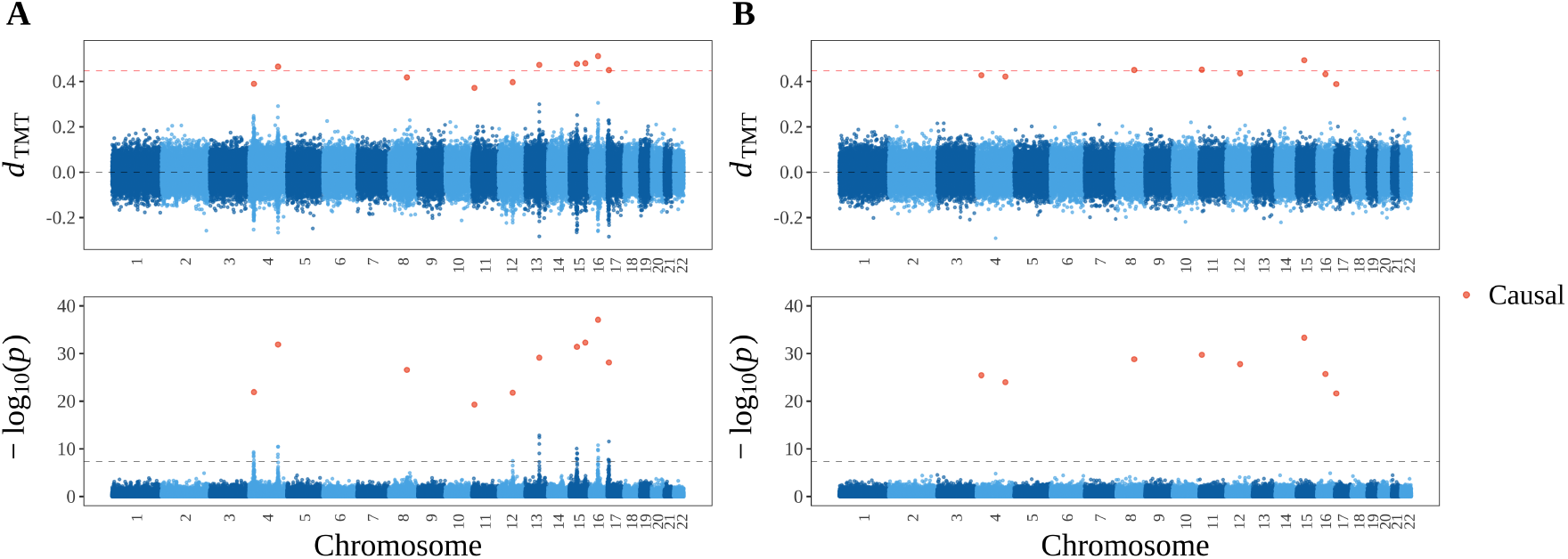
The genome-wide TMT profile. **(A)** Randomization linkage scenario. **(B)** Independent randomization scenario, where the genotypes from (A) were independently permuted to remove the randomization linkage. For both scenarios, the TMT profile is presented as *d*_TMT_ and − log_10_(*p*) at 100,000 SNPs across 22 chromosomes simulated by msprime. In the upper panel, the red dashed line is the true ACE at the causal loci and the black dashed line is for zero ACE at the non-causal loci. In the bottom panel, the gray dashed line is at *p*-value = 5 *×* 10^−8^, which is commonly used as a *p*-value threshold in GWAS.

### 3.4 TMT versus standard approaches in the presence of confounding

Regression approaches, such as the ordinary least square (OLS) and linear mixed model (LMM), identify genotypes that are associated with phenotypes while making no claims of causality. The OLS approach does not account for population structure, whereas the LMM approach does. In order to guarantee that association tests from these two approaches yield correct p-value and false positive rate calculations, both approaches make a nontrivial assumption of exogeneity, which means that the non-genetic variation has zero covariance with the genetic variation. Thus, when confounding factors are present that conflate these two sources of variation, these methods may yield inaccurate results, even at the level of association.

To assess the behavior of the TMT, OLS, and LMM approaches in the presence of exogeneity, for each child *j* ∈ [1 : *J*], we generated traits according to

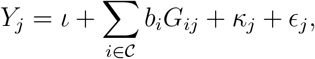

where 𝒞 is the set of causal SNPs and *κ*_*j*_ is confounded with a subset of non-causal (and unassociated) SNPs. Full details are in Section B.6. We implemented the TMT on a per marker basis to detect causal genotypes. We employed the LMM software GCTA [28] to detect associations. We also conducted simple linear regression between child trait and child genotypes on a per locus basis to test associations via OLS. In our simulation, the only SNPs that are associated with the trait are causal SNPs. Let *C* be the total number of causal SNPs and *U* the total number of non-causal confounded SNPs. Figure 6 displays the distribution of *p*-values at non-causal SNPs where we induced confounding and 9*U* other random non-causal and unconfounded SNPs. Since these 10*U* SNPs are non-causal (and unassociated with the trait), their *p*-values should be Uniform(0, 1) distributed. In our simulations, only the TMT results in Uniform(0, 1) distributed *p*-values of non-causal SNPs. Both LMM and OLS report significantly small *p*-values deviating from Uniform(0, 1) for non-causal SNPs.

**Figure 6:**
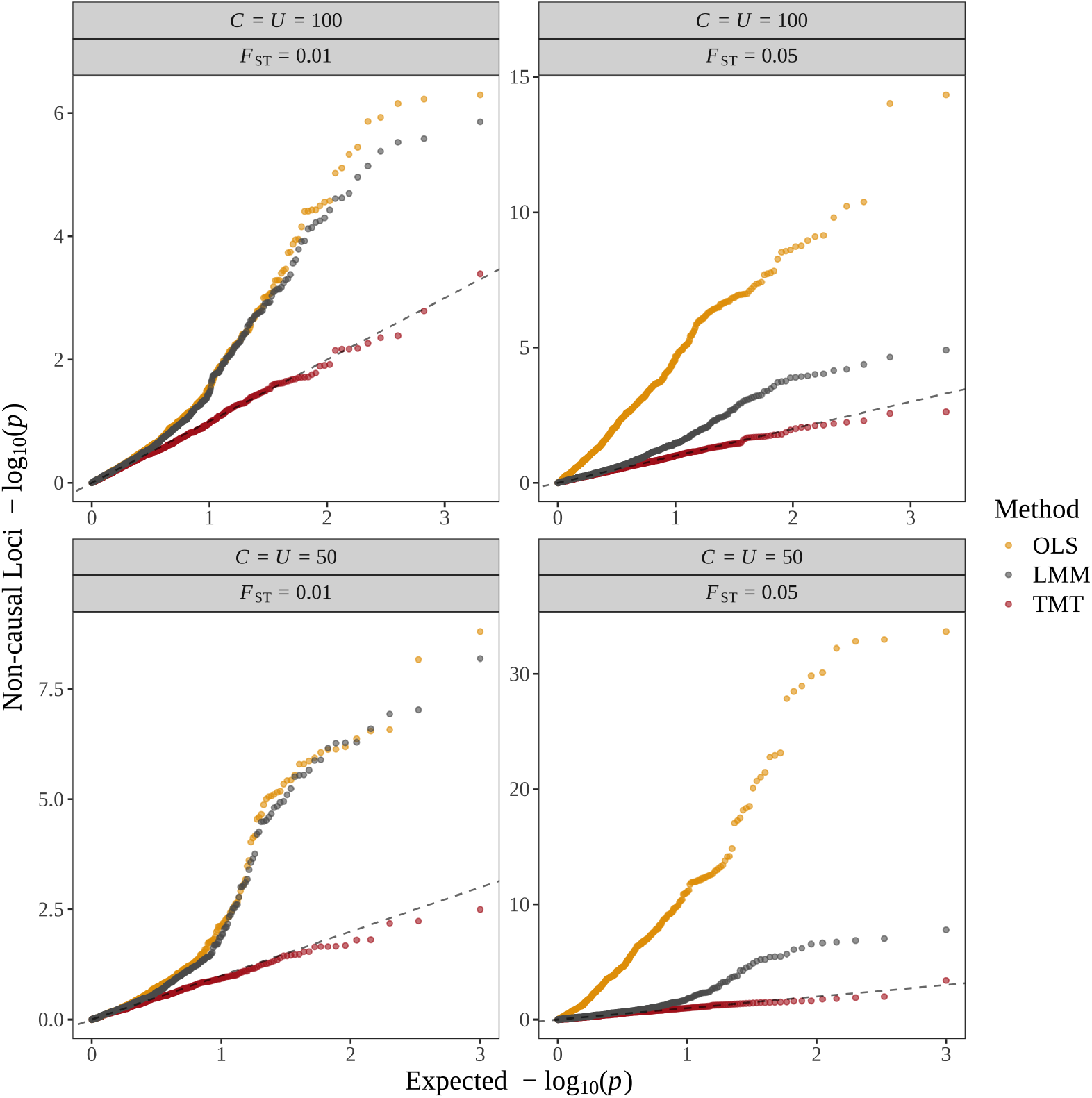
The distribution of − log_10_(*p*) at non-causal SNPs by the ordinary least square (OLS), the linear mixed model (LMM) and the TMT. The expected *p*-values on the x-axis are drawn from Uniform(0,1). Points are sorted by their quantiles.

### 3.5 Applying the TMT to large-scale association studies

Large-scale studies, such as the UK Biobank [14] and *All of Us* [15], involve hundreds of thousands of individuals. Many of the individuals in such large studies may be closely related. When studies contain closely related individuals (e.g., parent-child trios), researchers often retain only one of these individuals and exclude the others from the GWAS analysis. With the proposed TMT framework, one can instead set all closely related individuals aside and perform the GWAS on the remaining individuals. One can then follow up by validating the associations for causal effects on the related individuals using the TMT. Since the closely related individuals were not included in the initial GWAS analysis, one can even focus on a smaller set of the most significant SNPs from the GWAS analysis.

The UK Biobank has reported 107,162 related pairs (third degree or closer) [14] by examining kinship coefficients estimated with the software `KING` [47] and discordant homozygote rates. Following the criteria from `KING`, we extracted individual pairs with kinship estimates around 1/4 (between 2^−5*/*2^ and 2^−3*/*2^) as first degree relatives but not monozygotic twins. We further filtered out individual pairs with a discordant homozygote rate greater than 0.1, which are likely to be siblings. We classified the remaining individual pairs as parent-offspring pairs and identified families with at least one child and two parents (one male and one female). Through this procedure we identified 1,970 trio candidates, similar to previous studies [14, 48]. These sample sizes have reasonable power as shown in our simulations, especially on a well-selected subset of SNPs identified in a traditional GWAS as causal candidates. We expect a comparable or larger sample size of parent-child trios for the *All of Us* data since it has reported a much larger size of closely related participants, specifically 215,107 pairs (second degree or closer) [15].

As evidenced by the thousands of citations of the original TDT publications [20–22], trio studies play an important role in genetics. In dbGaP, trio studies with reasonable sample sizes are available, presenting both whole-genome sequencing data and phenotypes of interest. For example, the Pediatric Cardiac Genetics Consortium involves more than 5,000 trios, together with a disease status related to congenital heart disease; it aims to recruit 10,000 trios with affected and unaffected children [13, 49]. The total number of trios in these studies have similar sample sizes to our simulations. Therefore, in addition to the large-scale studies containing many trios, the proposed methods should be useful to apply to existing and forthcoming studies aimed at directly sampling trios.

## 4 Discussion

In studies where genetic trios (two parents and a child) are sampled from a population, it is possible to measure the transmission of genetic variants from parent to child. We have shown that by leveraging the inherent randomization in this process, it is possible to construct a potential outcomes model and inference method to rigorously test for causality from genetic variant to trait in the child. We proposed the transmission mean test (TMT) in this scenario for all common trait types – quantitative, count, or dichotomous. We showed that within our framework, the well-known transmission disequilibrium test (TDT), where trios are sampled based on a dichotomous affected trait status in the child, is also a test of causality.

We demonstrated both theoretically and empirically that the TMT and TDT are robust to a wide range of confounders, including population structure and the confounding of non-genetic variation with the genetic signal. Our theoretical analysis showed how genetic relatedness and structure of parents within and between trios do not affect the accuracy of our tests. More generally, our theoretical results clarify why the analysis of the concordance of genetic transmission and traits within and among many pedigrees is robust to common confounders. We showed how linkage disequilibrium and meiotic genetic linkage play roles in shared signal among neighboring SNPs.

The proposed framework is based on the potential outcomes model of causality in the context of randomized studies, which is the gold standard used in randomized controlled clinical trials required by the FDA [50]. Mendelian randomization is a method applied to standard GWAS where unrelated individuals from a population are sampled. Despite its name, Mendelian randomization does not observe the randomization process of genetic inheritance from parent to child. Rather, it is a method that implements instrumental variable regression using a genotype as the instrument. Mendelian randomization requires several assumptions, including no population structure [10]. A framework exists for the scenario where both parents and the child have phased genomes that utilizes a probabilistic independence definition of causality [12] and a computationally intensive simulation based null distribution [11], requiring Haldane’s model of recombination [51].

Given that standard GWAS can have very large sample sizes, one may consider the relevance of trio based studies. We noted that many trio studies of a reasonable sample size are present in dbGaP. We also discussed that large studies, such as the UK Biobank [14] and *All of Us* [15], contain many instances of first degree relatives, including full parent-child trios. A possible strategy is to set aside the first degree relatives and perform a standard GWAS on the remaining individuals to identify causal candidates based on associations. One can then employ the TMT on the related individuals on a reduced set of candidate SNPs to identify causal variants. Going beyond trios to more general first degree relatives, such as siblings, future work could develop a framework where one can first predict missing parental genotypes by Mendelian imputation [52] and then conduct the TMT based on this probabilistic imputation.

### Resources

Reproducible research documentation is available at https://github.com/StoreyLab/causal-trio. An R package `geneticTMT` that implements the proposed methods is available at https://github.com/StoreyLab/geneticTMT.

## Acknowledgments

This work was supported by US National Institute of Health grant R01 HG006448.

## Appendices

### A Theory

#### A.1 Proof of Lemma 1

Under the trait model in Equation (1),

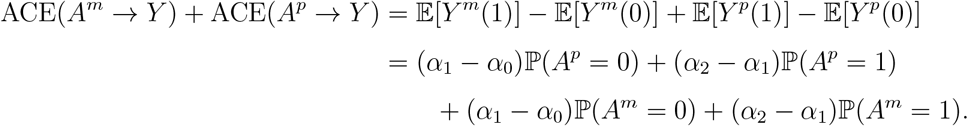

Assume 0 *<* ℙ(*A*^*m*^ = 1) *<* 1 and 0 *<* ℙ(*A*^*p*^ = 1) *<* 1. Under Assumption 1, either *α*_0_ ≤ *α*_1_ ≤ *α*_2_ or *α*_0_ ≥ *α*_1_ ≥ *α*_2_. Then ACE(*A*^*m*^ → *Y*) + ACE(*A*^*p*^ → *Y*) = 0 if and only if *α*_0_≠ *α*_1_ or *α*_1_ ≠ *α*_2_, which means ACE(*G* → *Y*)≠ 0 by Definition 2.

#### A.2 Proof of Lemma 2

Here we show (*Y* ^*m*^(0), *Y* ^*m*^(1) **╨** *A*^*m*^|*Z*^*m*^ = 1). The proof for (*Y* ^*p*^(0), *Y* ^*p*^(1)) **╨** *A*^*p*^|*Z*^*p*^ = 1 is the same. We first show that *A*^*p*^ **╨** *A*^*m*^|*Z*^*m*^ = 1 through several calculations.

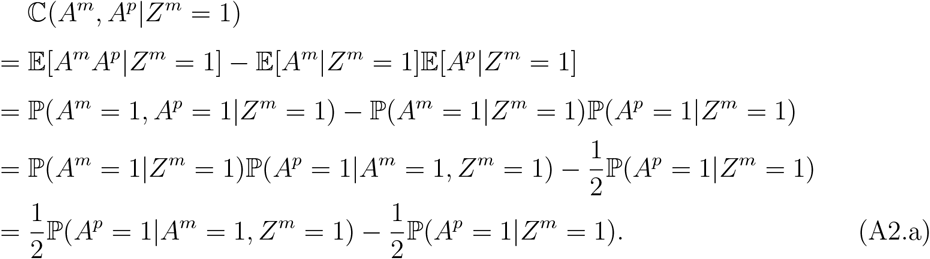

By the Law of Total Covariance,

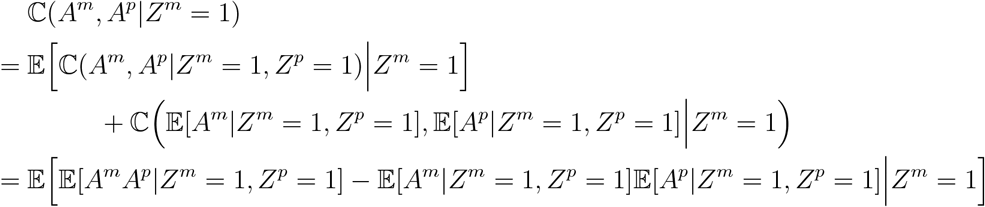

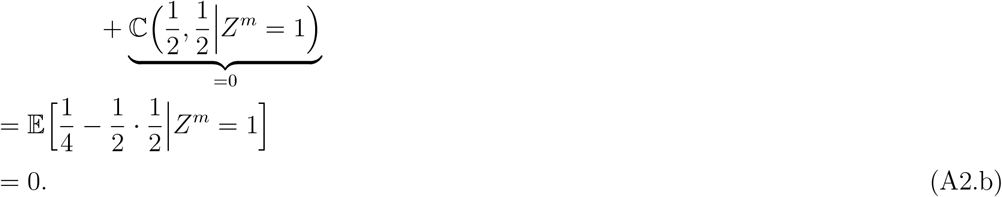

Since Equation (A2.a) and Equation (A2.b) are equal, this implies ℙ(*A*^*p*^ = 1|*A*^*m*^ = 1, *Z*^*m*^ = 1) − ℙ(*A*^*p*^ = 1|*Z*^*m*^ = 1) = 0. Similarly, ℙ(*A*^*p*^ = *b*|*A*^*m*^ = *a, Z*^*m*^ = 1) = ℙ(*A*^*p*^ = *b*|*Z*^*m*^ = 1) for *a, b* ∈ *{*0, 1*}*. So

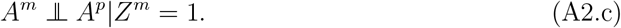

By Assumption 2,

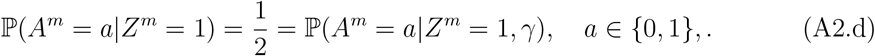

Under the trait model in Equation (1), *Y* ^*m*^(0) and *Y* ^*m*^(1) are functions of *A*^*p*^ and *γ*. By Equation (A2.c) and Equation (A2.d), we have shown that *A*^*p*^ and *γ* are independent of *A*^*m*^ conditional on *Z*^*m*^ = 1. Thus, (*Y* ^*m*^(0), *Y* ^*m*^(1)) |*Z*^*m*^ = 1 and *A*^*m*^|*Z*^*m*^ = 1 are independent.

#### A.3 Proof of Lemma 3

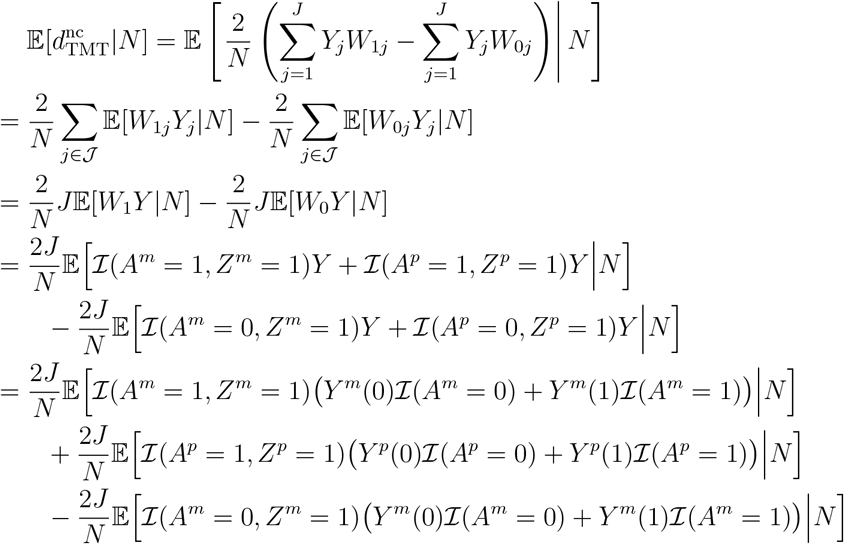

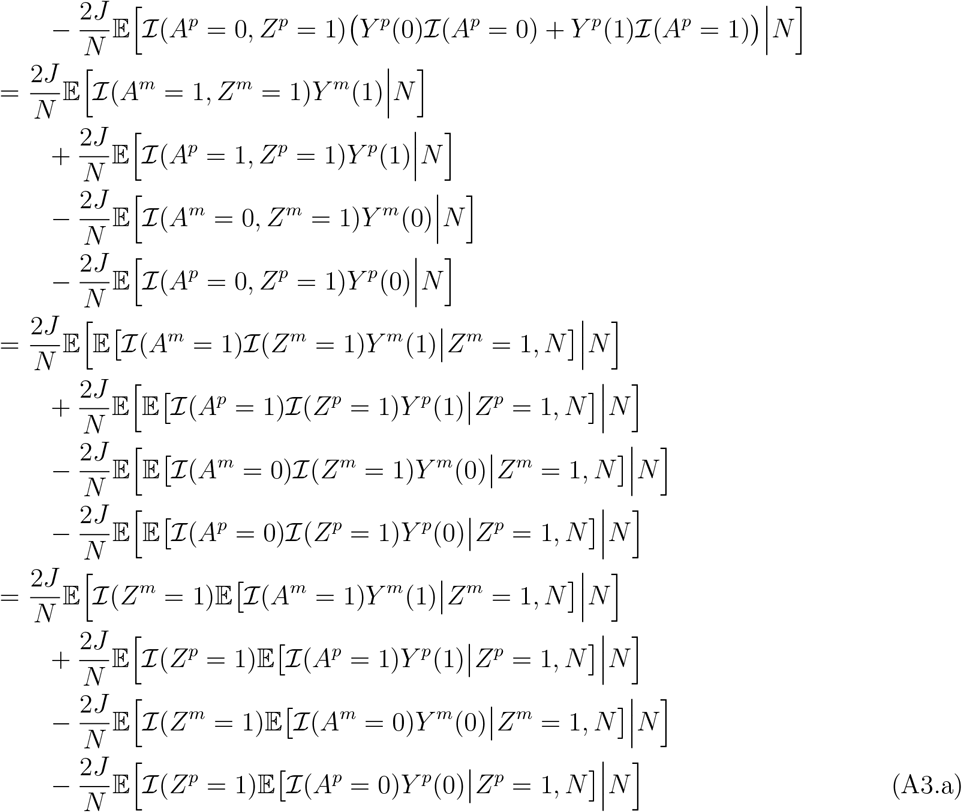

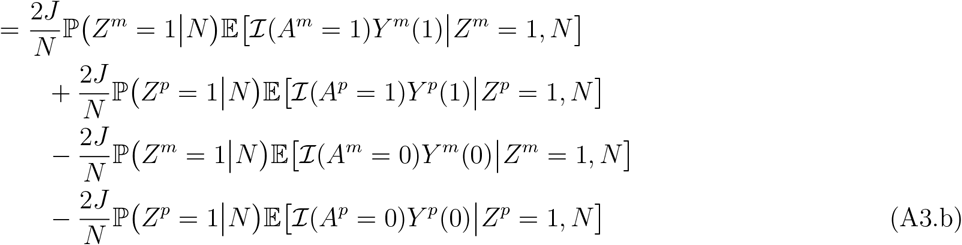

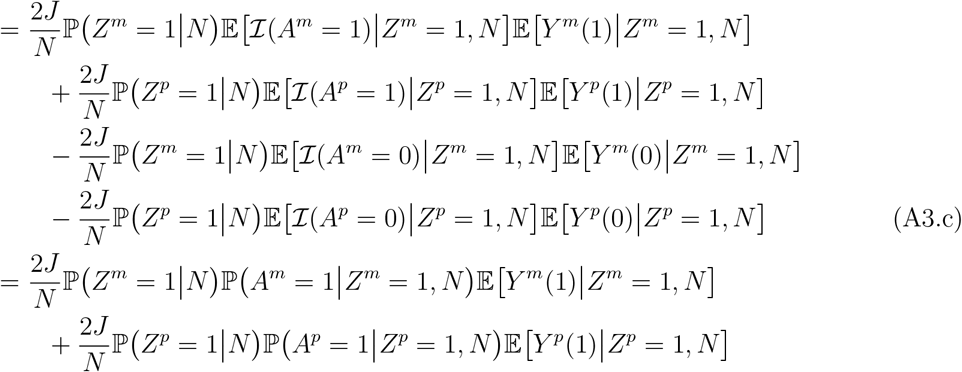

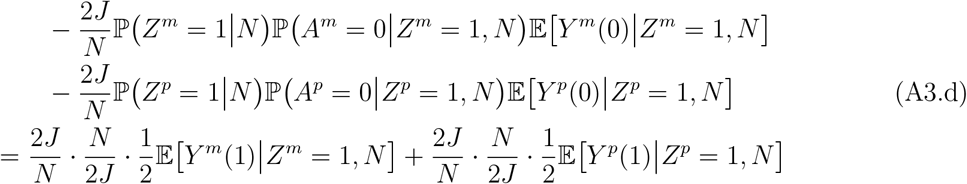

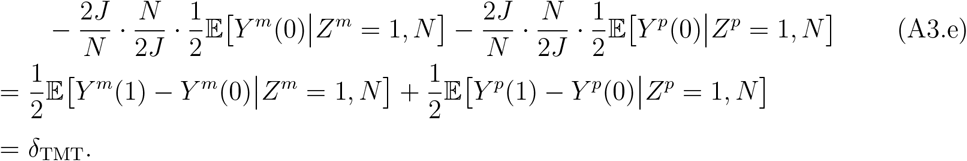

Equation (A3.a) is equal to Equation (A3.b) by the Law of Total Probability. Equation (A3.b) is equal to Equation (A3.c) by Lemma 2 in that *Y* (0), *Y* (1) **╨** *A Z* = 1. Equation (A3.d) is equal to Equation (A3.e) since 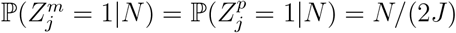, which follows because all parents are exchangeable and there are 2*J* parents.

#### A.4 Proof of Lemma 4 Part A

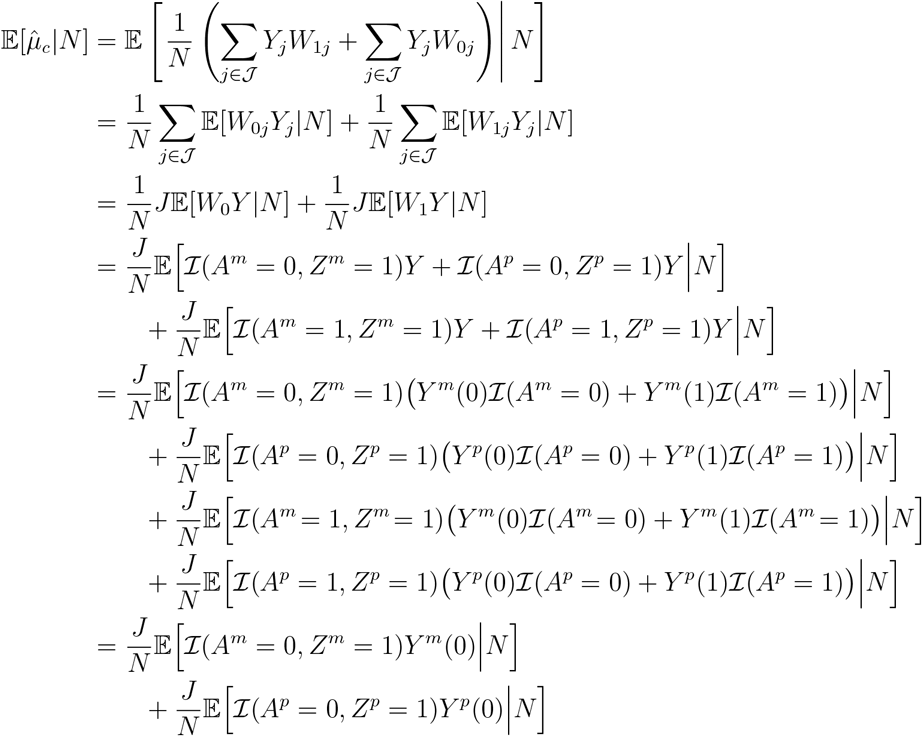

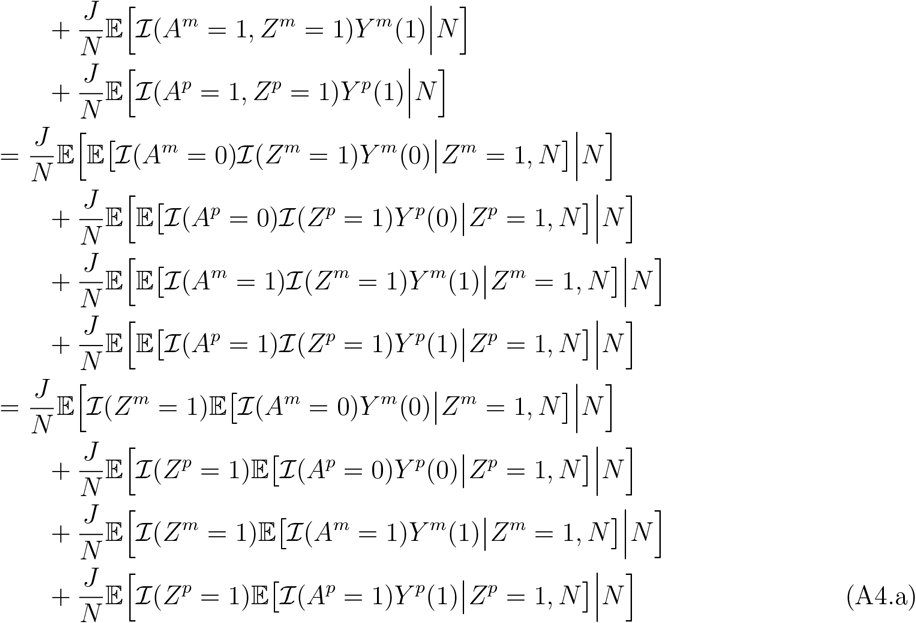

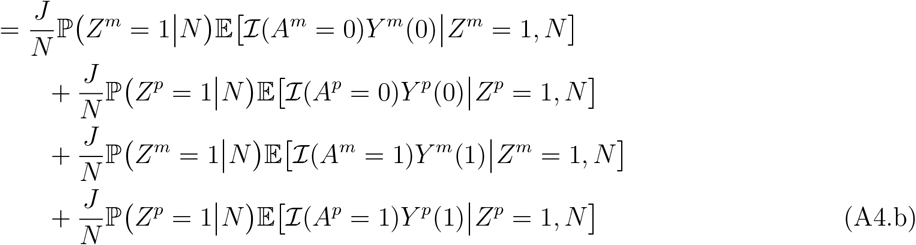

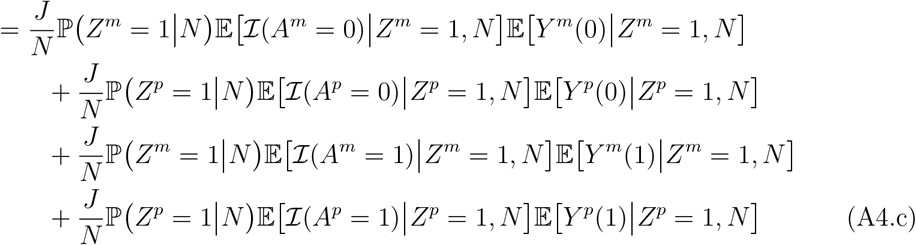

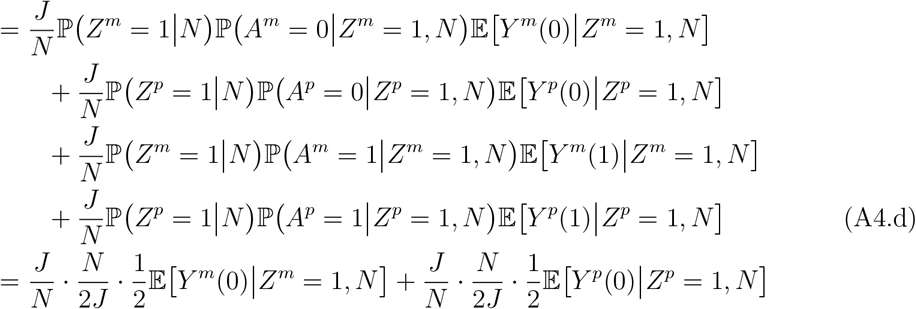

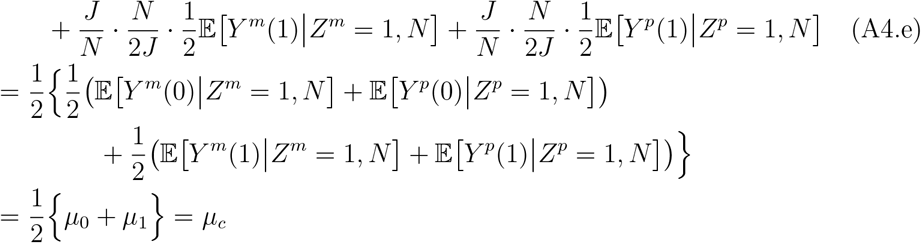

This completes the proof for Lemma 4. Equation (A4.a) is equal to Equation (A4.b) by the Law of Total Probability. Equation (A4.b) is equal to Equation (A4.c) by Lemma 2 in that *Y* (0), *Y* (1) **╨** *A Z* = 1. Equation (A4.d) is equal to Equation (A4.e) because 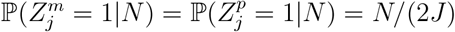,, which follows because all parents are exchangeable and there are 2*J* parents.

#### A.5 Proof of Lemma 4 Part B

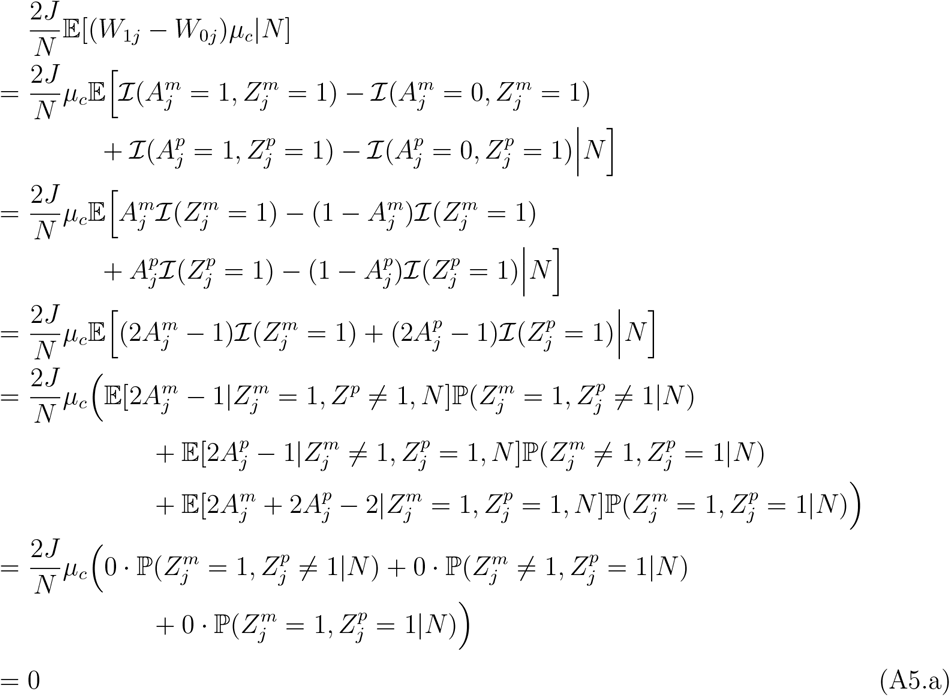

To show 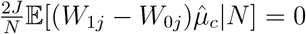, first define

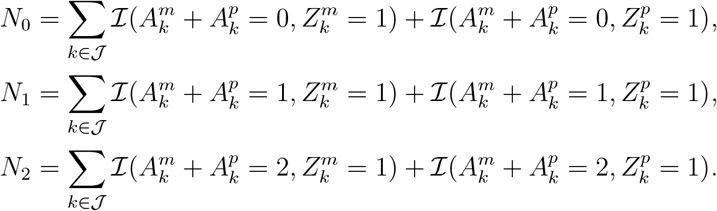

Notice that *N*_0_ + *N*_1_ + *N*_2_ = *N*. Recall that

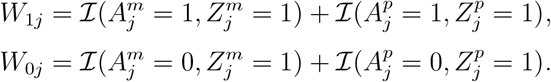

Then

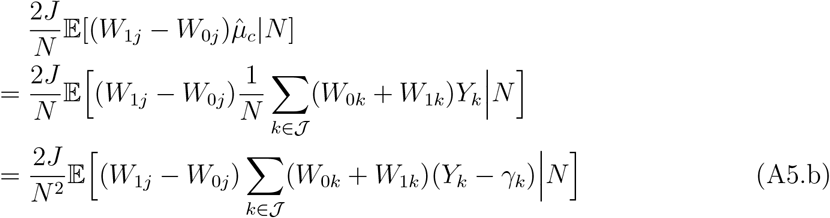

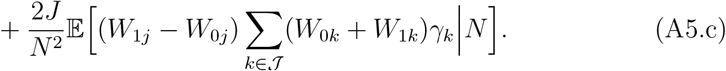

For Equation (A5.b), it equals

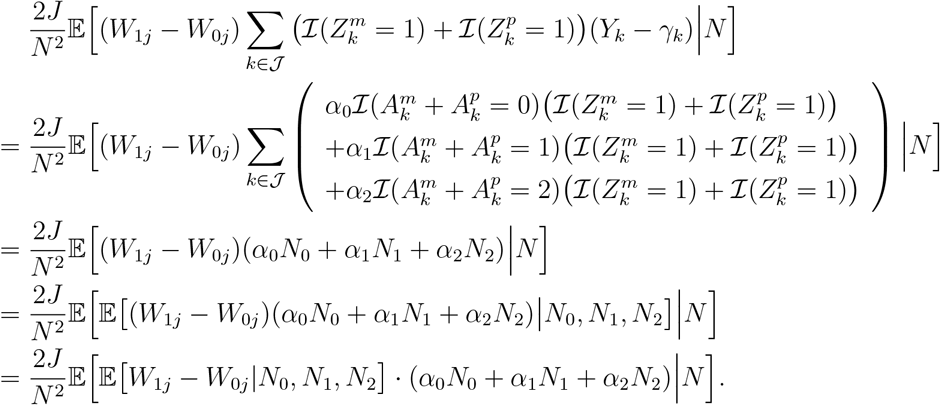

Note that

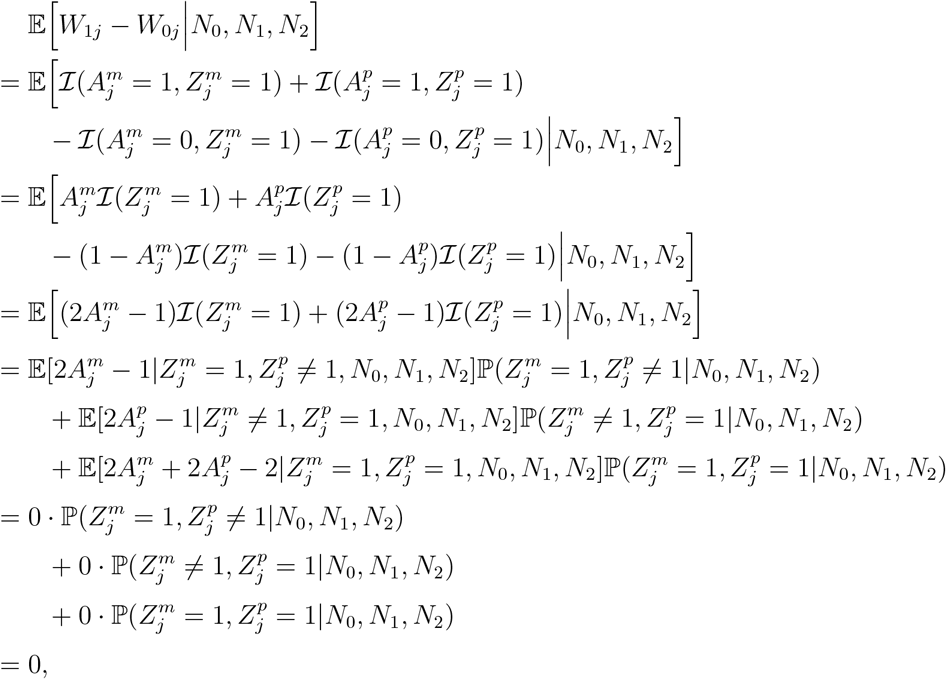

and thus,

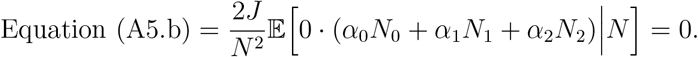

For Equation (A5.c), it equals

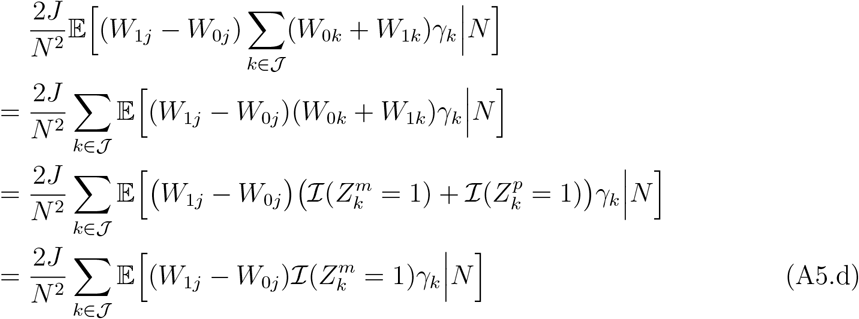

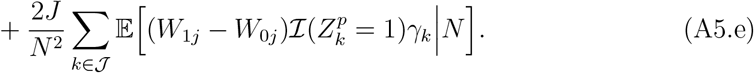

For Equation (A5.d), notice that

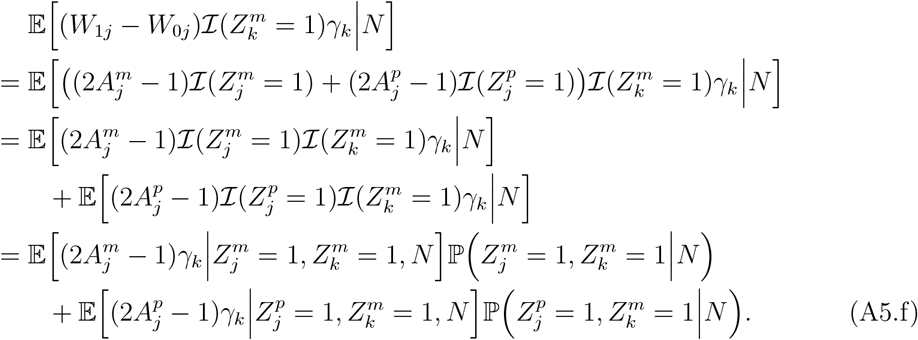

By the Law of Segregation,

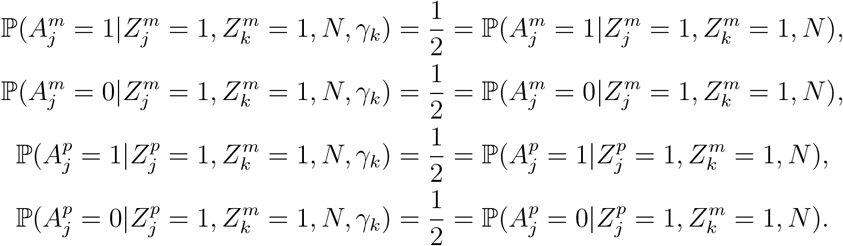

Then Equation (A5.f) equals

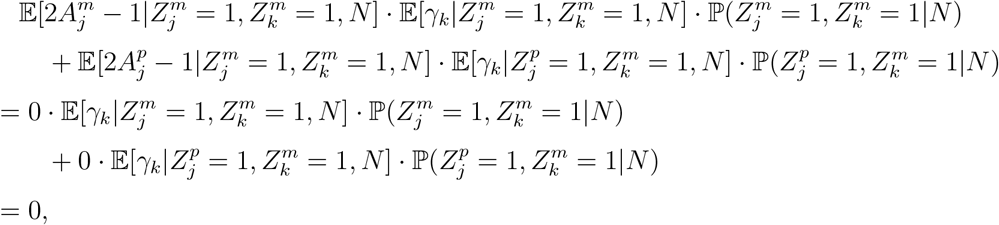

and thus Equation (A5.d) = 0. Similarly, Equation (A5.e) = 0. Then Equation (A5.c) = Equation (A5.d) + Equation (A5.e) = 0. Therefore, Equation (6) in Lemma 4 holds in that

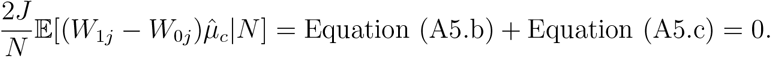

#### A.6 Proof of Lemma 5

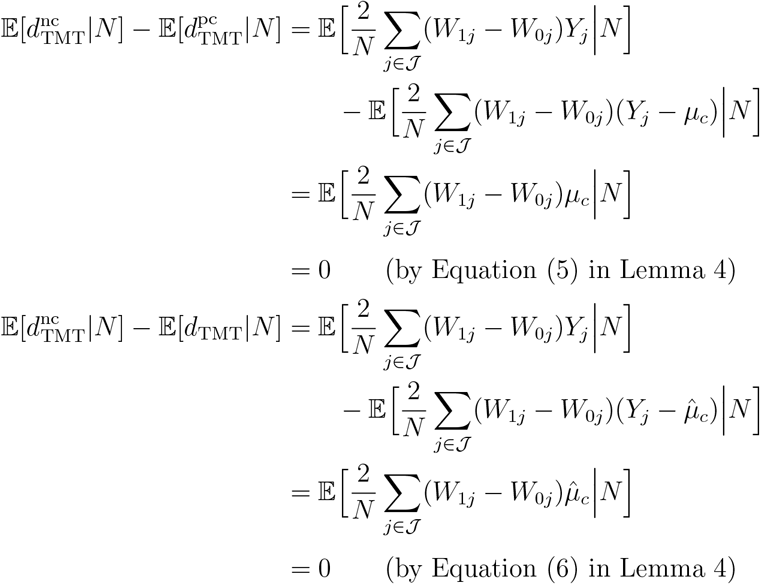

Thus, 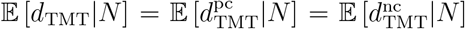. By Lemma 3, 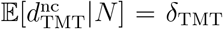. Therefore, 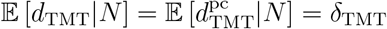.

#### A.7 Proof of Lemma 7

For the parental transmitted allele, *A*, under Assumption 3,

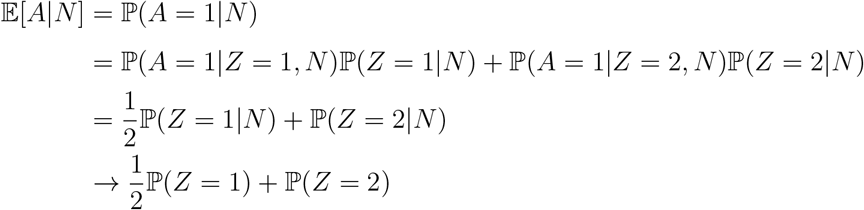

with probability 1 when *J* → ∞. Notice that

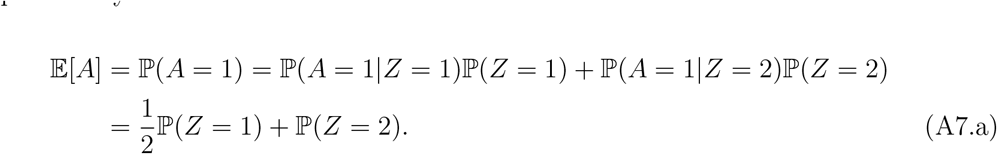

Therefore, 𝔼[*A*|*N*] → 𝔼[*A*] with probability 1 when *J* → ∞. Also,

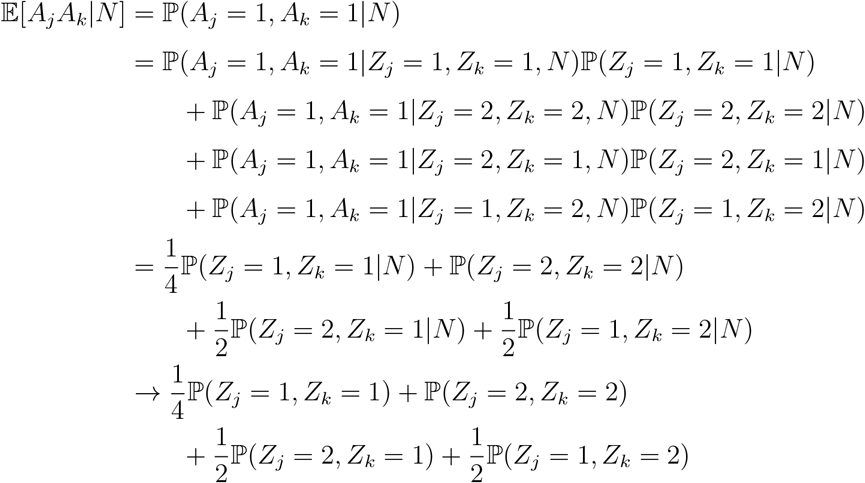

with probability 1 when *J* → ∞. Notice that

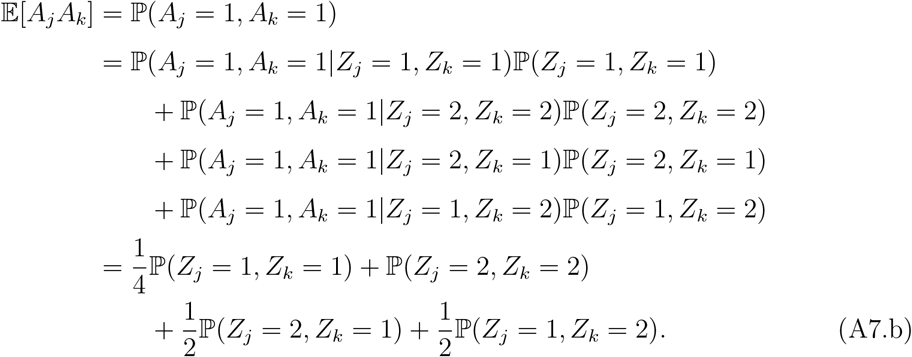

Then 𝔼[*A*_*j*_*A*_*k*_|*N*] → 𝔼[*A*_*j*_*A*_*k*_] with probability 1 as *J* → ∞. So far we have shown:

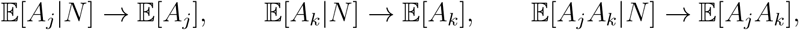

with probability 1 as *J* → ∞. Then by Slutsky’s Theorem,

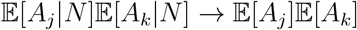

as *J* → ∞. Together, we derive

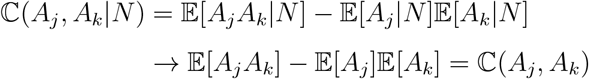

as *J* → ∞. Now we show 𝕔(*A*_*j*_, *A*_*k*_) ≥ 0. By Equation (A7.a) and Equation (A7.b),

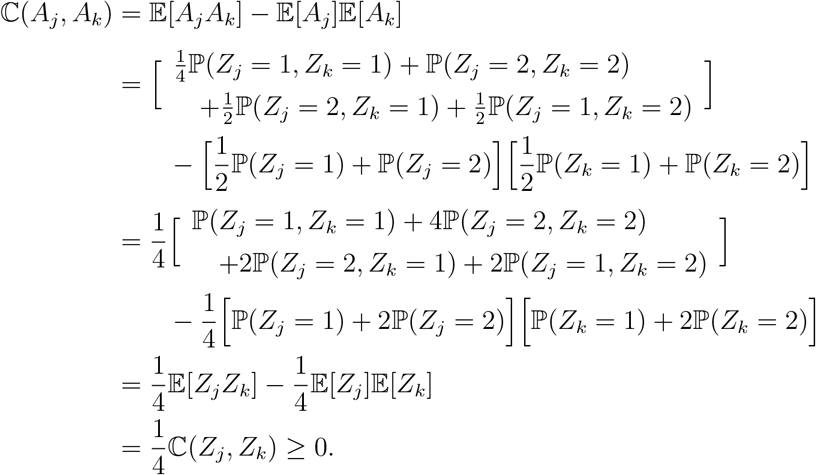

#### A.8 Aroof of Lemma 8

Let 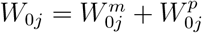 and 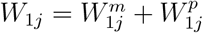 where

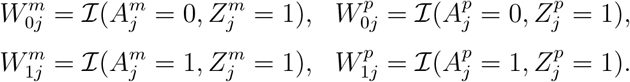

Then

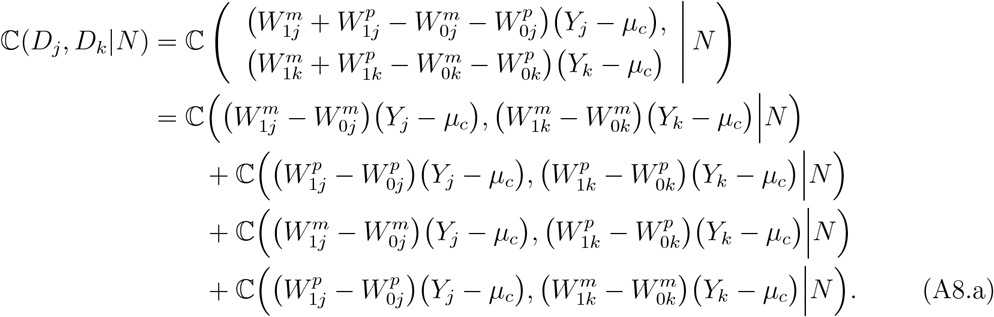

Equation (A8.a) involves four covariances whose calculations follow the same algebra. For the first covariance

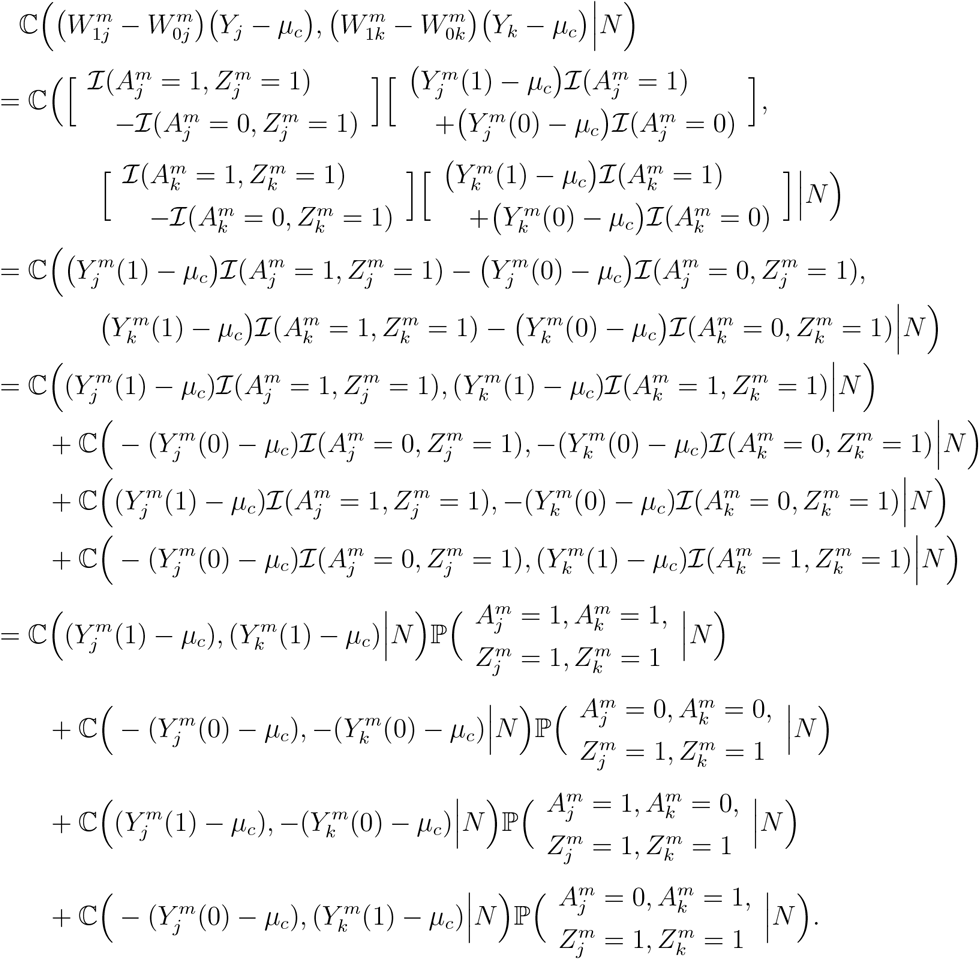

Notice that

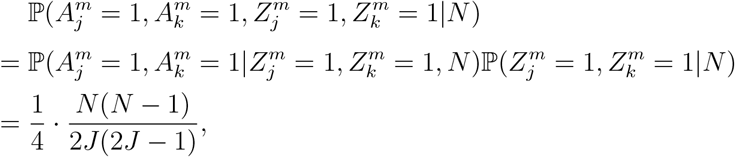

and also

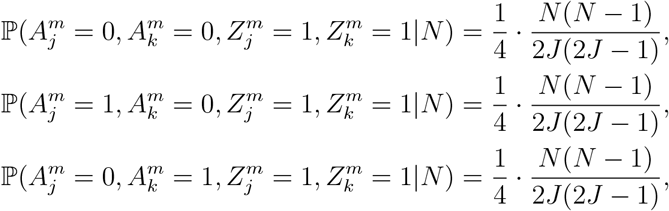

then

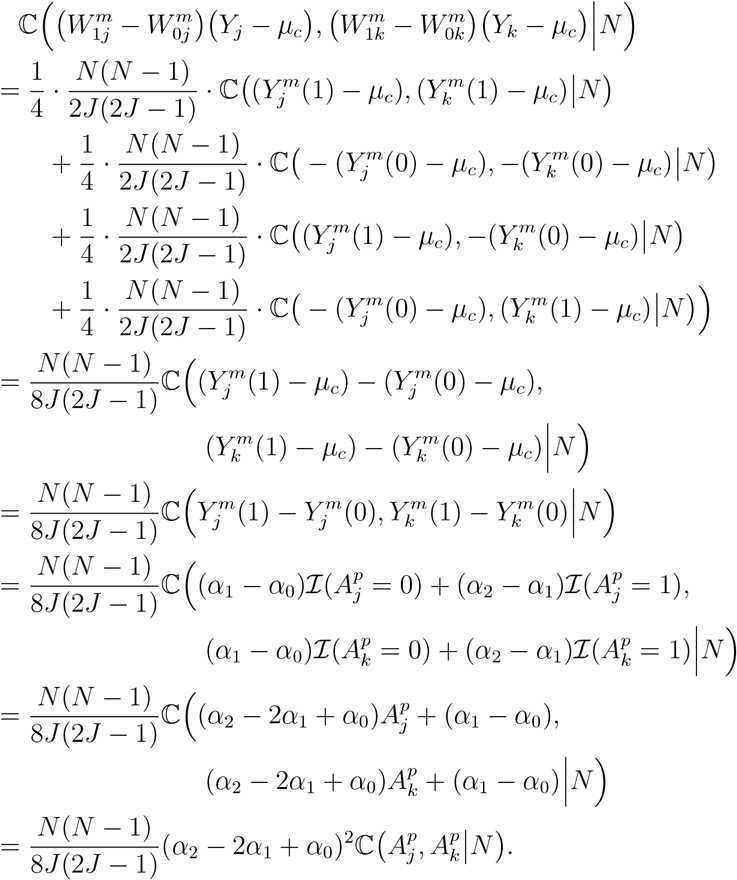

By Lemma 7, when 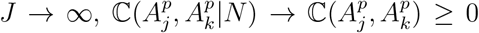 Also by Assumption 3, let ℙ(*Z* = 1) = *ω*, then ℙ(*Z* = 1|*N*) = *N/*2*J* → *ω* with probability 1 when *J* → ∞. Therefore,

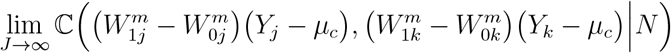

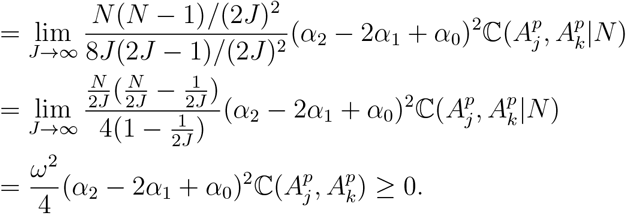

One can follow the same approach to calculate non-negative asymptotic values for the other three covariances in Equation (A8.a). Adding up these four covariances and noting they all have non-negative asymptotic values, implies

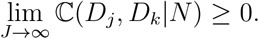

The equality, lim_*J*→∞_ 𝕔(*D*_*j*_, *D*_*k*_|*N*) = 0, holds for the following scenarios:

A. Under a true null hypothesis where *α*_0_ = *α*_1_ = *α*_2_ so that (*α*_2_ − 2*α*_1_ + *α*_0_) = 0.
B. Under a true alternative hypothesis where *α*_1_ −*α*_0_ = *α*_2_ −*α*_1_ so that (*α*_2_ −2*α*_1_ +*α*_0_) = 0. This setting is equivalent to an additive genetic effect model.
C. The genotypes of the *j*th and the *k*th children have zero covariance.

#### A.9 Proof of Lemma 9

Let *D*_*j*_ = (*W*_1*j*_ − *W*_0*j*_)(*Y*_*j*_ − *µ*_*c*_) and 𝒯 = {𝒯_0_, 𝒯_1_, 𝒯_00_, 𝒯_11_}. Then 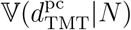 equals

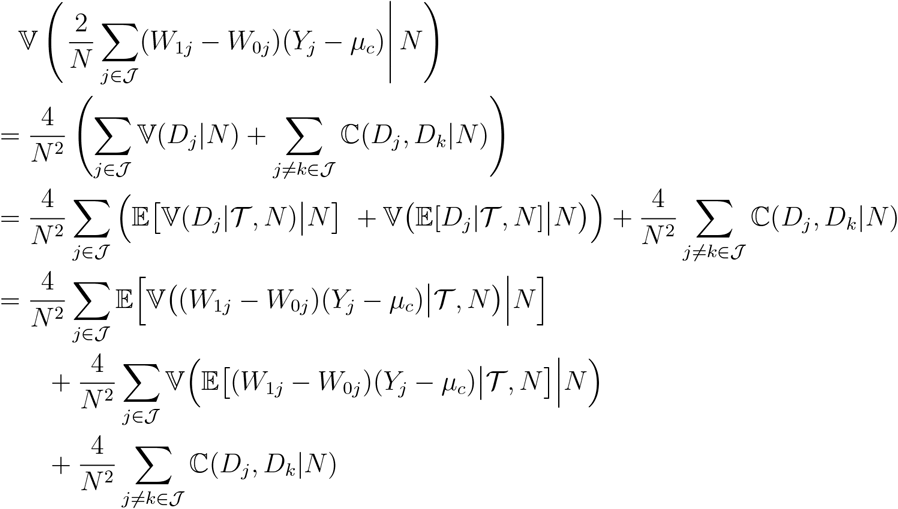

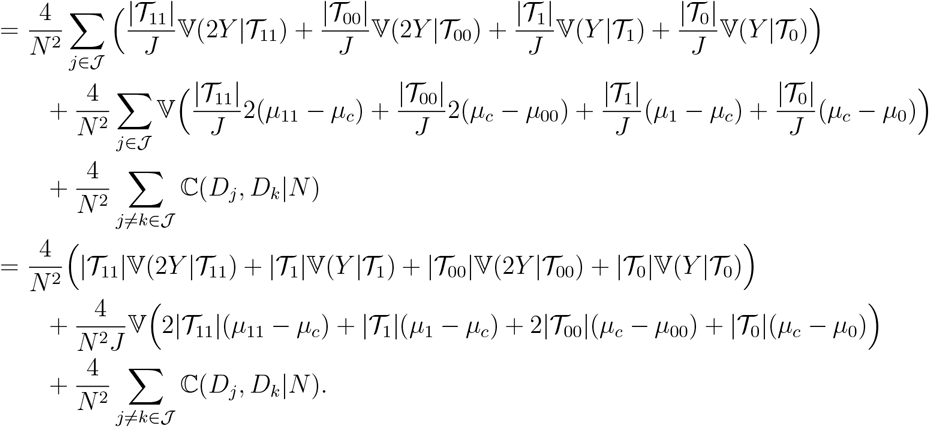

Let

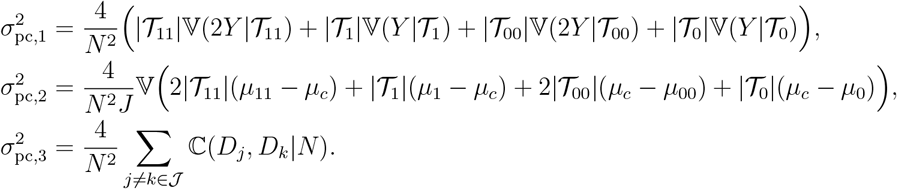

By Lemma 6, 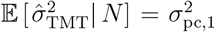. When the null hypothesis is true, then *µ*_11_ = *µ*_1_ = *µ*_*c*_ = *µ*_0_ = *µ*_00_ so that 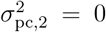. Under a true alternative hypothesis,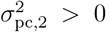. By Lemma 8, when the null hypothesis is true then 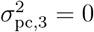. When the alternative hypothesis is true 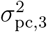 converges almost surely to a non-negative value as *J* → ∞. Since 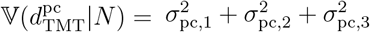, it follows that:

A. under null, 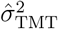 is an unbiased estimator for 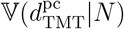
B. under alternative, 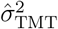 underestimates 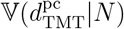 as *J* → ∞.

As an aside,

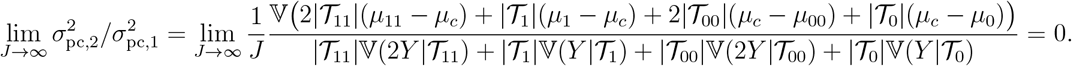

#### A.10 Proof of Lemma 10

It is the case that

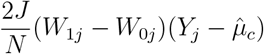

for *j* ∈ 𝒥 are exchangeable with each term having finite first absolute moment. Given these properties, by Lemma 4 and Equation 2.2 of ref. [53] implies

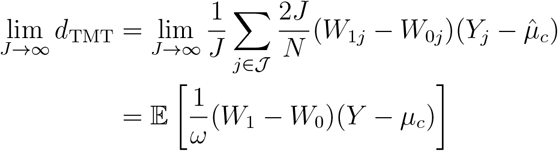

almost surely. Similarly,

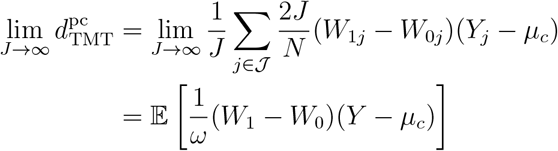

almost surely. Therefore,

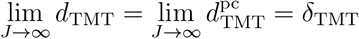

almost surely. Together with Lemma 5 where 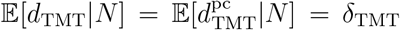, by the Continuous Mapping Theorem,

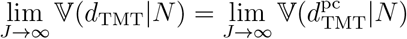

almost surely. In other words, 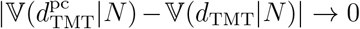 with probability 1 as *J* → ∞.

#### A.11 Proof of Lemma 11

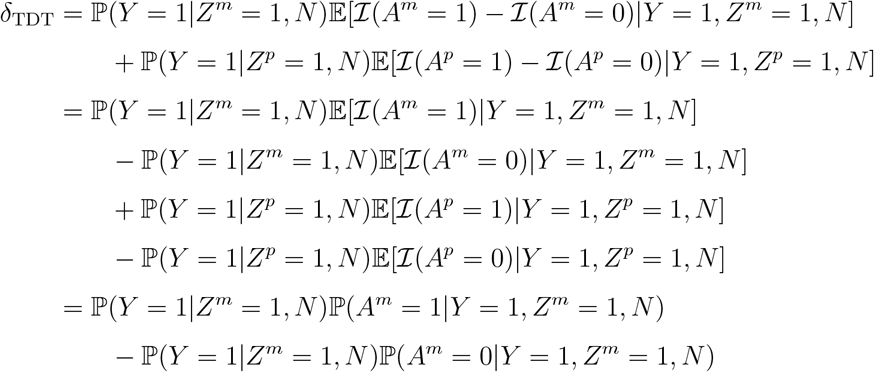

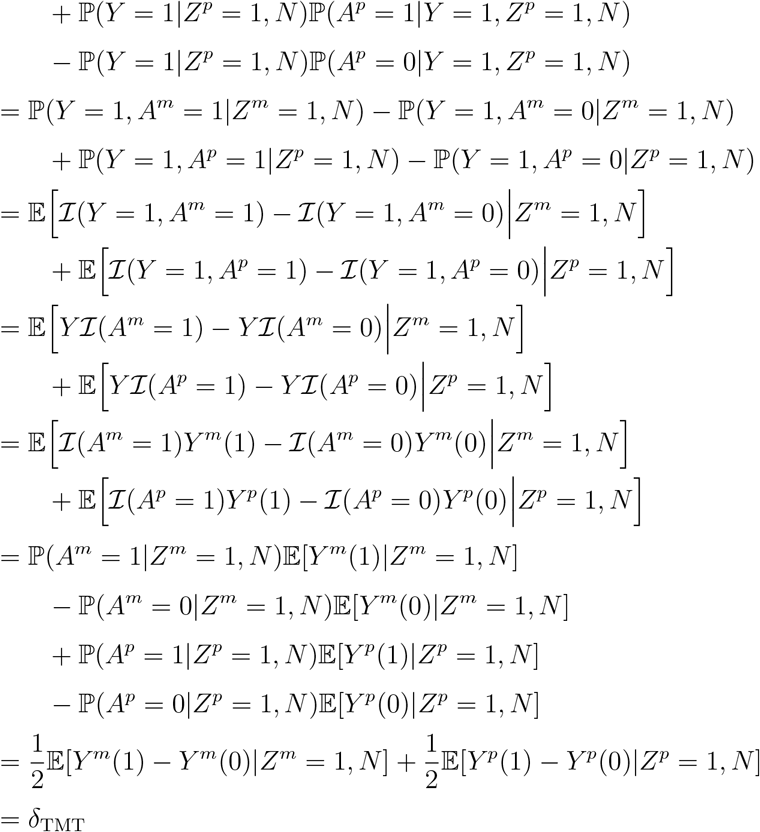

#### A.12 Proof of Lemma 12

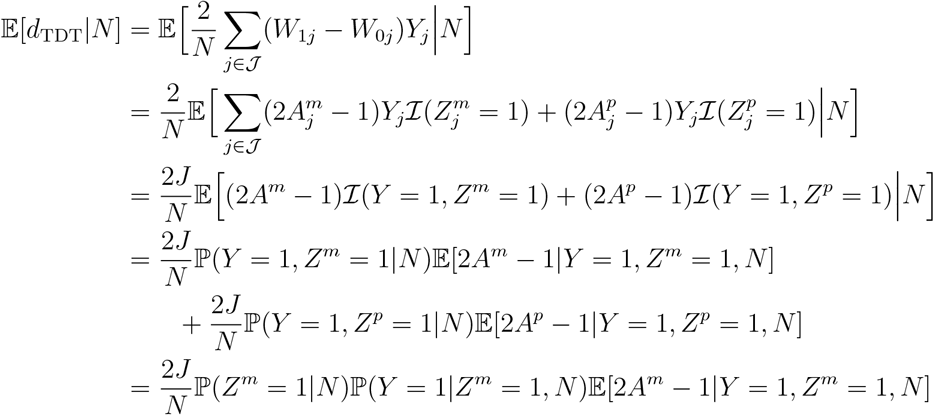

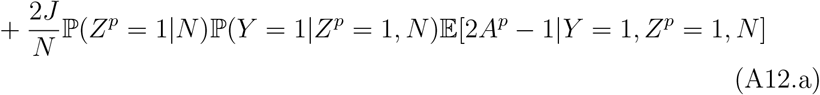

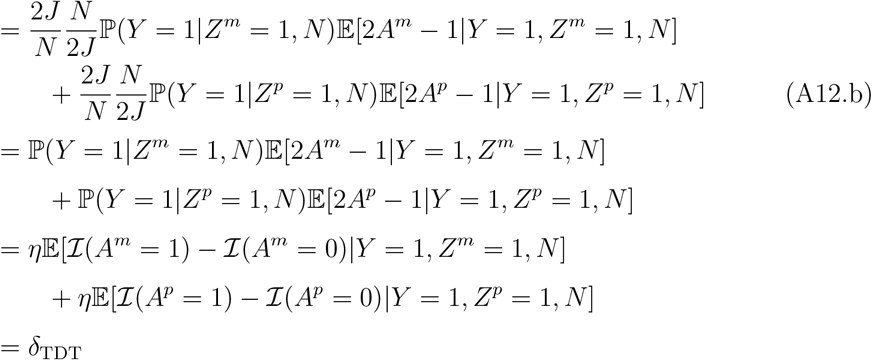

Line (A12.a) to line (A12.b) is due to the fact that 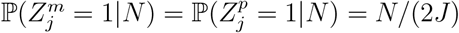.

#### A.13 Proof[U1] of Lemma 13

By Theorem 1, *δ*_TMT_ ≠ 0 at marker *d* if and only if ACE(*G*_*d*_ → *Y*)≠ 0. By Theorem 2, *δ*_TDT_≠ 0 at marker *d* if and only if ACE(*G*_*d*_ → *Y*) ≠ 0. Then by Lemma 1, ACE(*G*_*d*_ →*Y*)≠ 0 if and only if 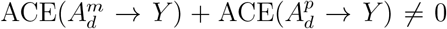, which is equivalent to either 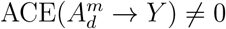 or 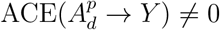 under Assumption 1. This means either 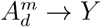 or 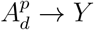 according to Definition 3. This further implies that either 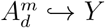 or 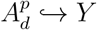 by Definition 11 since marker *d* is in complete linkage with itself, which means 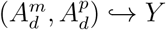 by Definition 12.

### B Simulations

#### B.1 Simulating trio genotypes

Both parental genotypes matrices ***Z***^*m*^ and ***Z***^*p*^ were sampled from a structured population based on a standard admixture model [54, 55] with *K* = 4 admixed populations. We configured this population to have *F*_ST_ = 0.2 by utilizing the `bnpsd` R package [33, 56]. We simulated the ancestral allele frequencies from the Uniform(0.1, 0.9), which is an option in the bnpsd R package. To visualize the population structure, we presented the co-ancestry coefficients between individuals from the structured population in Figure S2.

#### B.2 Simulating quantitative trait

We first generated a non-genetic factor associated with the population structure. Let ***E*** = {*E*_*j*_} be the random non-genetic factor. We adopted the admixture proportion ***q*** = {*q*_*ju*_} from Section B.1 to simulate *E*_*j*_ as

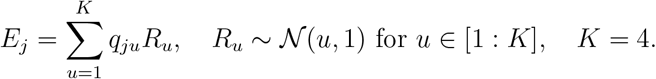

Let 𝒞 be the loci indices set for all causal SNPs. We followed Equation (13) to simulate the child’s trait ***Y*** = {*Y*_*j*_} as

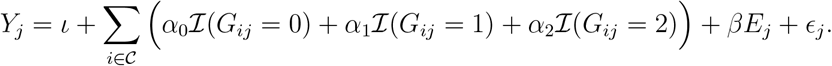

To show that the above equation satisfies the trait model in Equation (1), at the *c*th causal SNP (*c ∈* 𝒞), rewrite the equation as

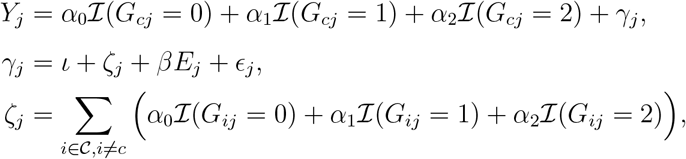

where the first line satisfies Equation (1). We considered three sets of (*α*_0_, *α*_1_, *α*_2_) as follows

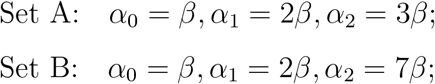

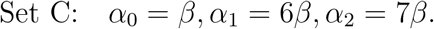

Then the ratio (*α*_2_ *− α*_1_)*/*(*α*_1_ *− α*_0_) is 1, 5 and 1*/*5 for the above Set A, B and C. Given the heritability 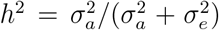 where we set 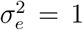, we simulated *β* such that 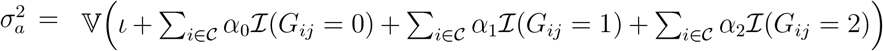, achieving the desired

#### B.3 Permutation test

This is done by permuting the observed trait for *B* times and generate a *B*-length vector of the TMT statistic. In each round of permutation, let 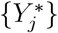 be the permuted trait values, calculate 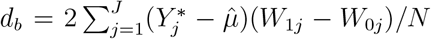 Then calculate *p*-value = 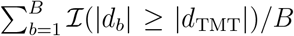 Without additional notes, we set the permutation times *B* = 1, 000.

#### B.4 Simulating dichotomous trait

We started from Equation (14) to generate a continuous latent variable ***L*** = {*L*_*j*_}. Let Ψ(·) be the probit function which is the inverse of standard Normal cumulative distribution function. We standardized *L*_*j*_ as 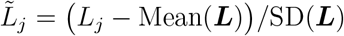. G iven a disease pre valence *λ*, 0 *< λ <* 1, we generated the dichotomous trait value by 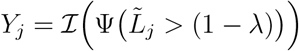

#### B.5 Simulating genetic linkage

For the parental genotypes, we used the software `msprime` (version 1.0) [43] and followed the American Admixture model [44] to simulate 5,000 pairs of parents from the admixed Americans popoulation with 100,000 SNPs across 22 chromosomes per individual, each chromosome with 2 haplotypes. Our simulation parameters for `msprime` are the same as parameters listed in the supporting information for [44]. We used the R package `popkin` [33] to calculate the *F*_ST_ of the parental genotypes, which is around 0.18. We set the total number of SNPs in each chromosome proportional to the corresponding chromosome length in Human Genome Assembly GRCh38.p14. We set the mutation rate per generation so that the allele frequency is between 0.05 and 0.95. We set the recombination rate so that the level of LD matches previous findings in human genome [45, 46]. Within each family, for each chromosome, we randomly drew a haplotype per parental side and merge these two haplotypes as the child genotypes.

#### B.6 Simulating confounding effects

We followed Section B.1 to simulate parental genotypes ***Z***^*m*^, ***Z***^*p*^ and child genotypes ***G*** with *F*_ST_ = 0.01, 0.05 for the underlying population. Let ***Q*** be the random variable that has confounding effects on the relationship between ***G*** and *Y*. We considered the confounding effects from parental genotypes via 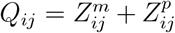. Let 𝒞 be the set of all causal SNPs and *U* the set of non-causal confounding SNPs. We denote the size of these two sets by *C* = | 𝒞 | and *U* = |𝒰 |. We generated the child trait by

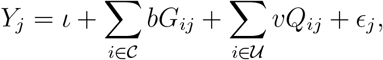

where we drew *ϵ*_*j*_ from Normal(0, 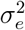) and we set 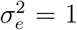, *ι* = 100. The coefficients *b* and *v* were determined such that 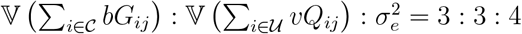

### C Supplementary Figures

**Figure S1:**
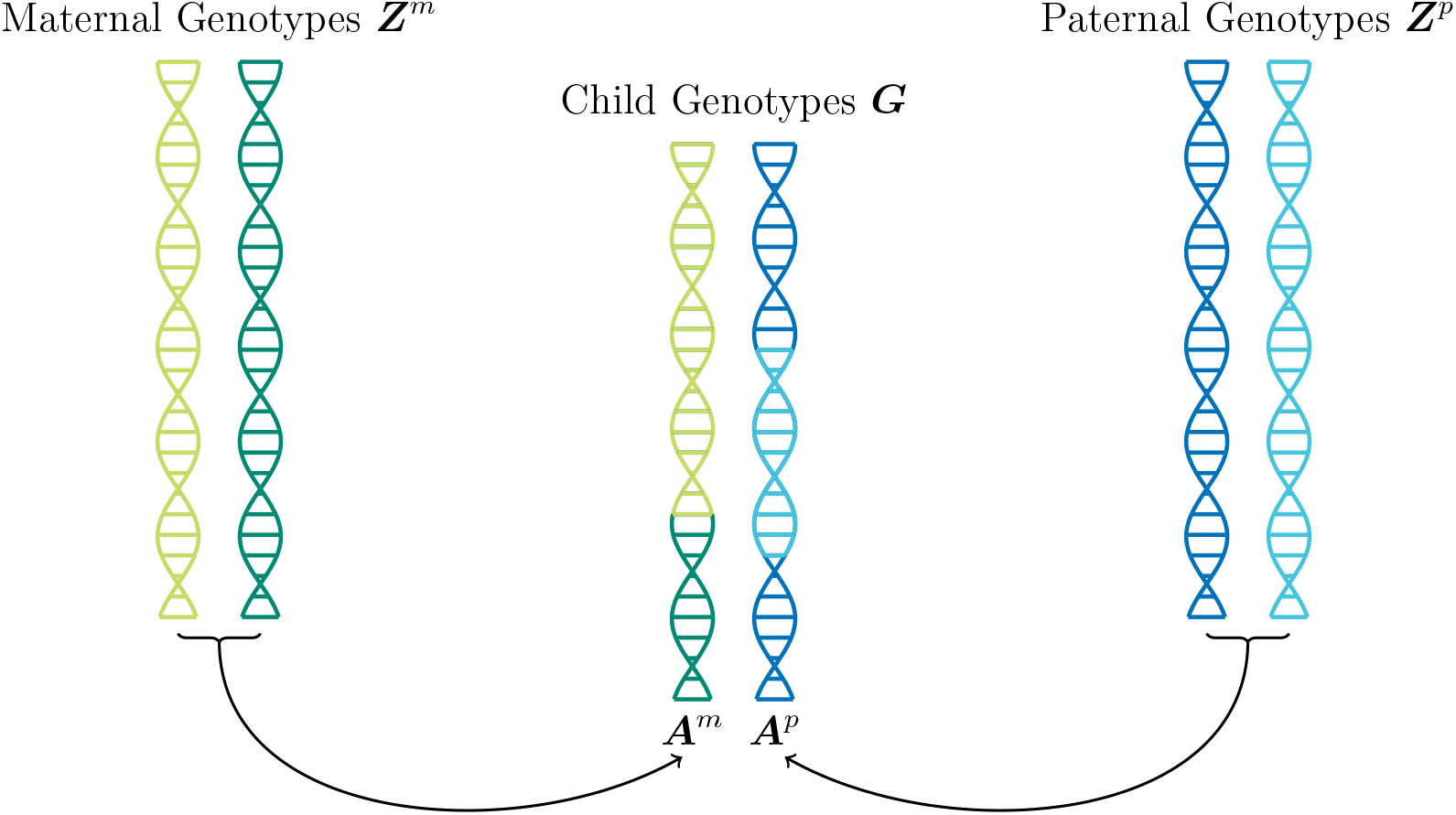
Schematic of a typical set of trio data and the meiosis process. The transmission of alleles from heterozygous parents to the child motivate the randomized experiments to identify causal relationship between the child’s phenotypes and genotypes. Recombination events may occur, marked as the adjacent dark and light colored pieces for the child’s genotypes.

**Figure S2:**
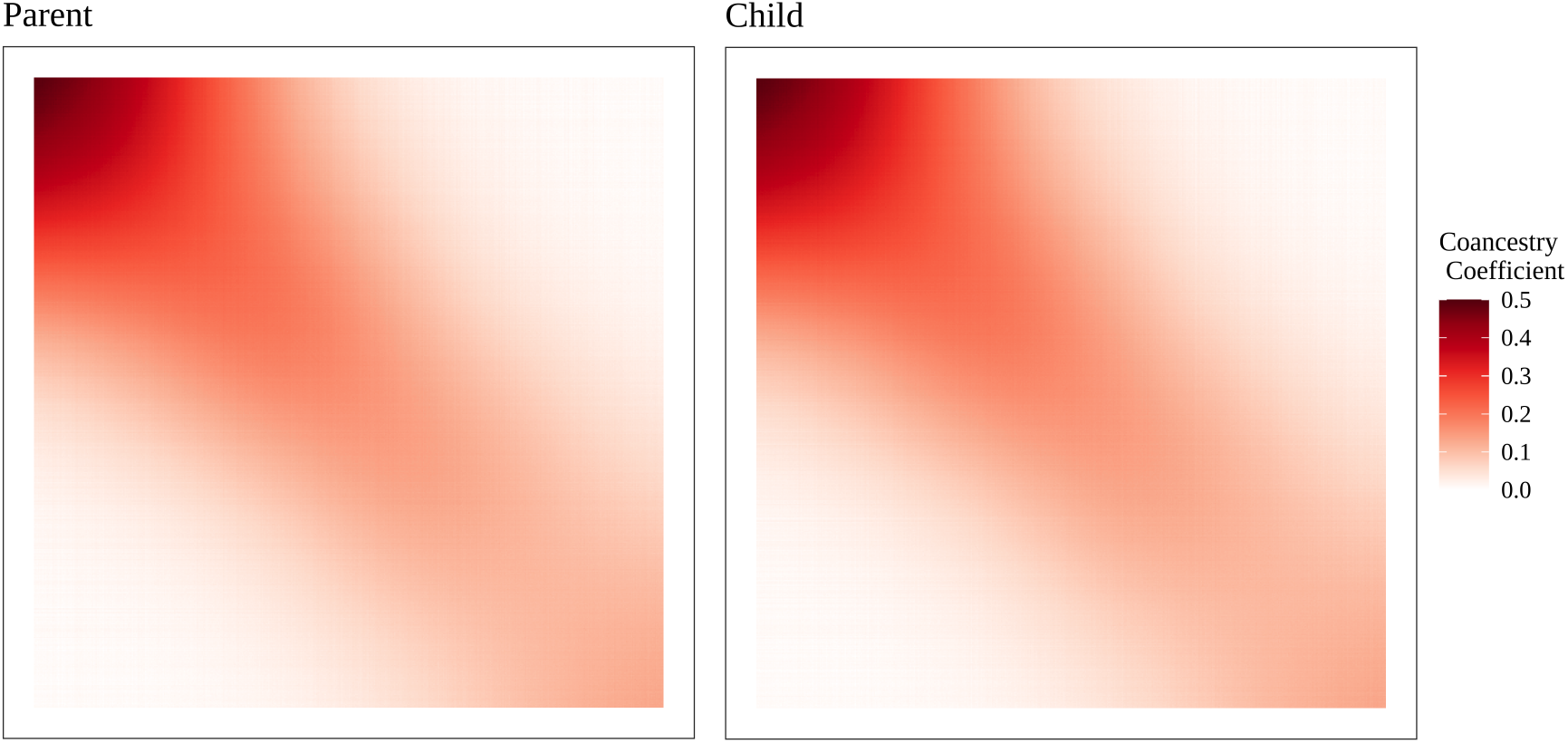
The co-ancestry coefficients between individuals among the structured population. Along both x-axis and y-axis lined up the individuals, mothers for the left panel and children for the right panel. Each entry is a pair-wise coancestry coefficient calculated by `popkin` [33] for a simulated sample of 500 trios randomly drawn from the structured population. Here we presented a sample of 500 trios due to the limited figure size. Simulations throughout the paper with larger sample sizes share the same pattern of population structure and the same variability of coancestry coefficients with the overall *F*_*ST*_ = 0.2.

**Figure S3:**
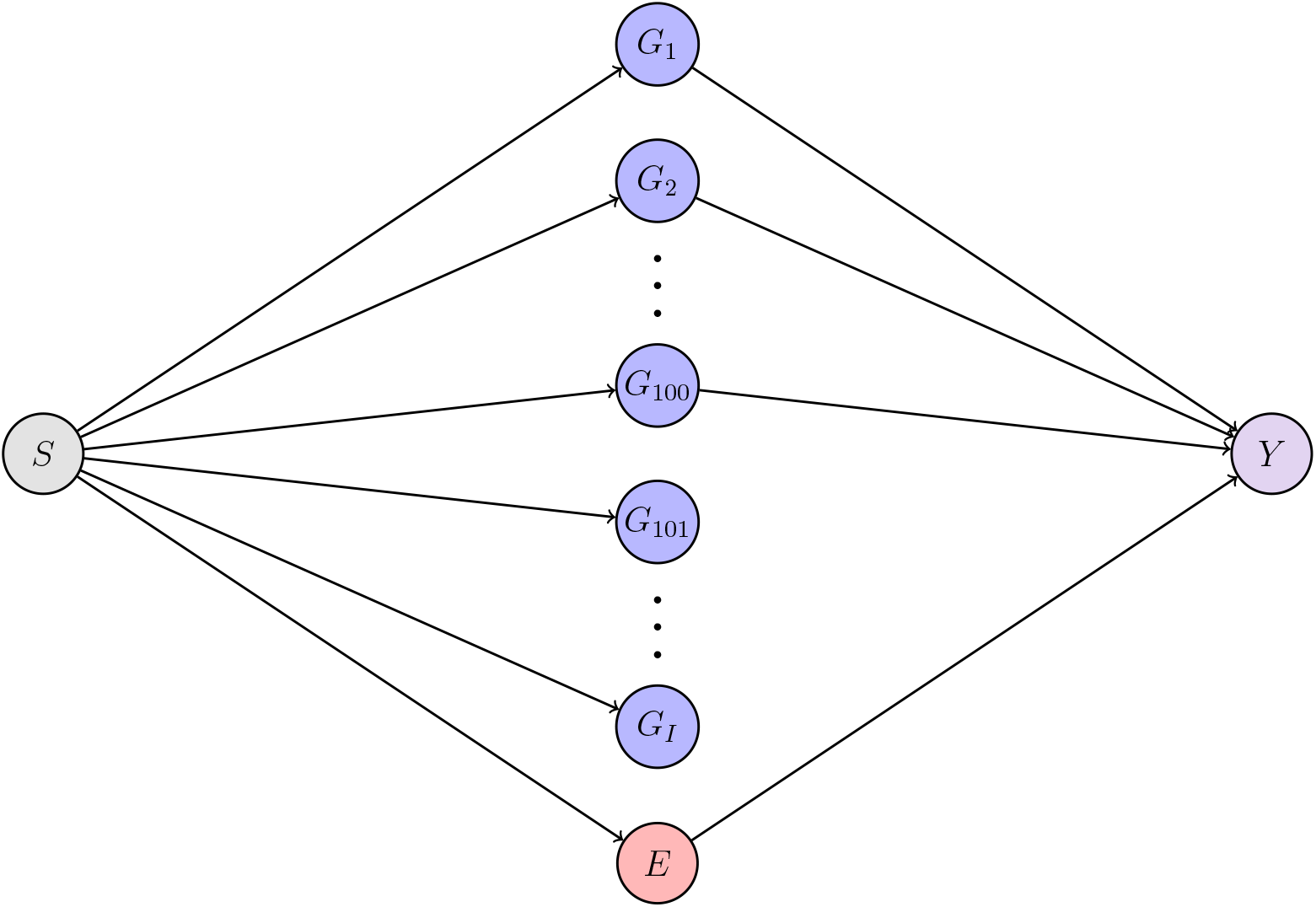
Schematic of the data simulation model. The phenotype *Y* is a function of 100 directly causal loci (*G*_1_, *G*_2_, …, *G*_100_) and a non-genetic factor *E*. The remaining loci (*G*_101_, *G*_102_, …, *G*_*I*_) are not causal for *Y*. All non-genetic and genetic variables (*E, G*_1_, *G*_2_, …, *G*_*I*_) are probabilistically dependent according to the population structure, represented by the variable *S*.

**Figure S4:**
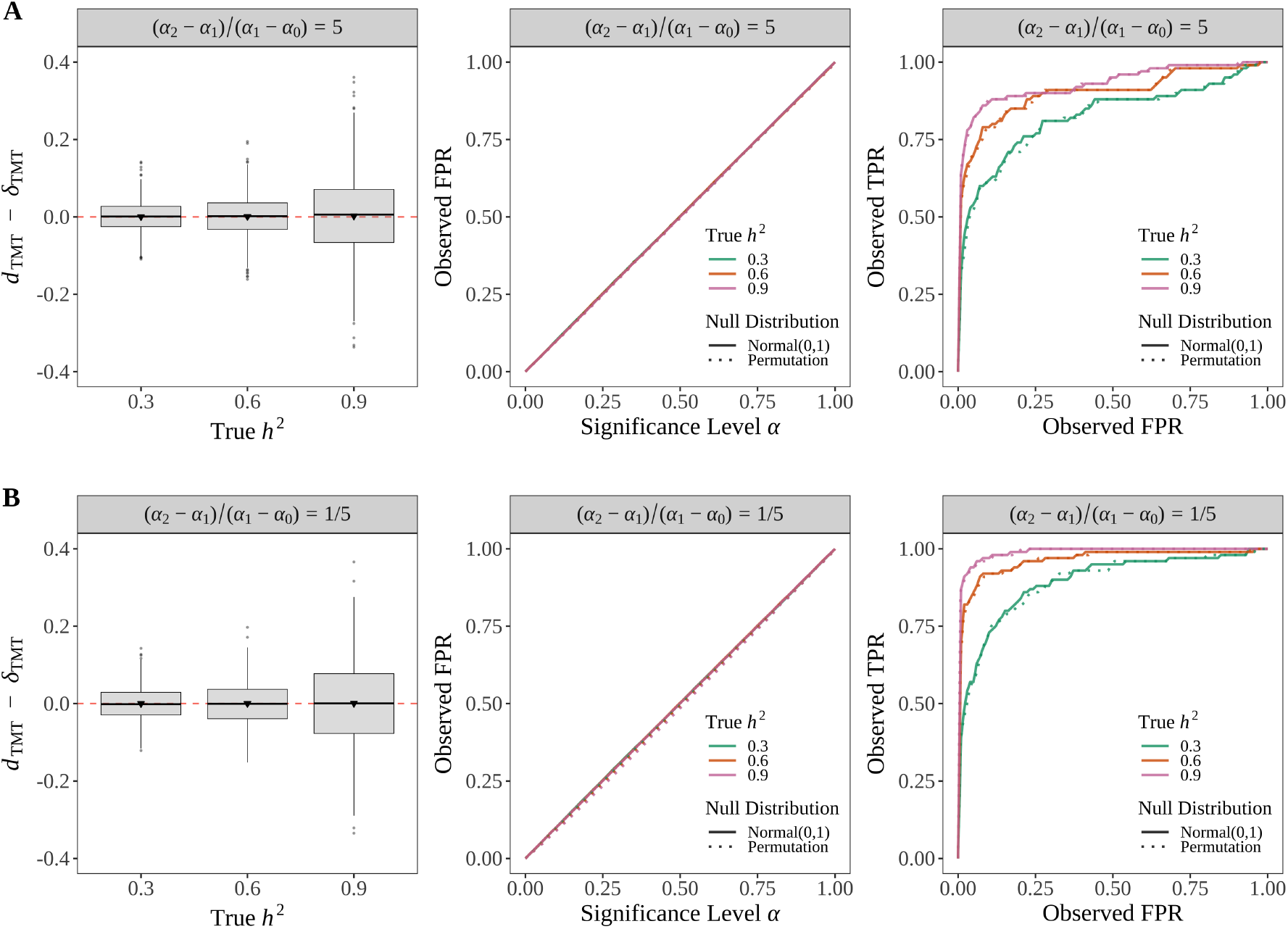
The TMT delivers unbiased causal effect estimand *d*_TMT_, controls FPR at the desired significance level, and has reliable ROC curve across various levels of heritability *h*^2^. **(A)** Simulation results for (*α*_2_ *− α*_1_)*/*(*α*_1_ *− α*_0_) = 5. **(B)** Simulation results for (*α*_2_ *− α*_1_)*/*(*α*_1_ *− α*_0_) = 1*/*5. We followed Section B.1 to draw 5, 000 trios from an admixed population (*F*_*ST*_ = 0.2) with 100, 000 SNPs and 100 causal loci per individual. We simulated quantitative traits by following Section B.2.

**Figure S5:**
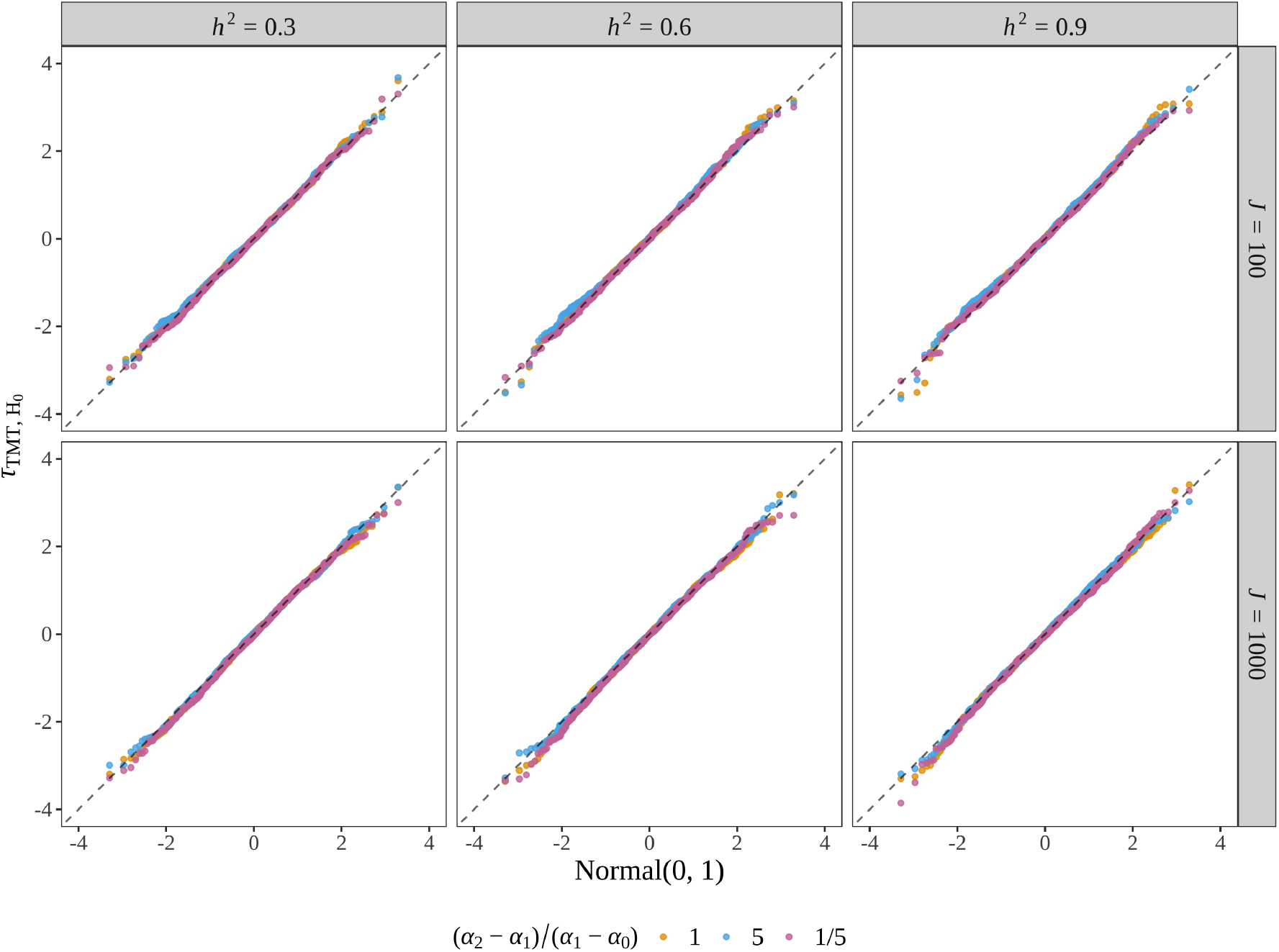
The Q-Q plot for the statistic *τ*_TMT_ under null. We followed Section B.1 to draw 100 and 1, 000 trios from an admixed population (*F*_*ST*_ = 0.2) with 100, 000 SNPs and 100 causal loci per individual. The desired *h*^2^ is 0.5. We simulated quantitative traits by following Section B.2. For a randomly chosen causal locus, we conducted 1,000 permutations on trait values and calculated the statistic *τ*_TMT_ per permutation to generate the observed null.

**Figure S6:**
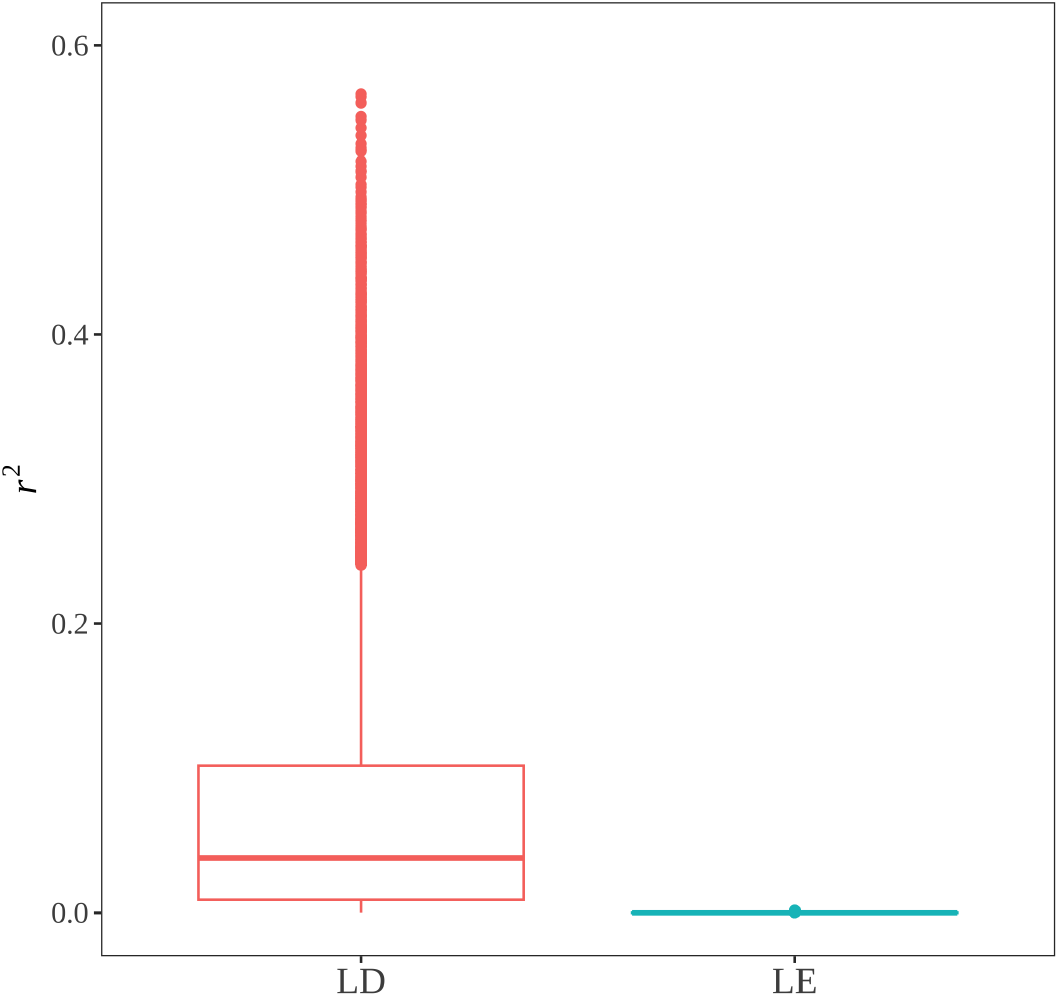
The distribution of *r*^2^ between adjacent SNPs. The *r*^2^ values are shown for all pairs of adjacent SNPs for the simulated linkage-disequilibrium (LD) scenario and the linkage-equilibrium (LE) scenario after permutation.

**Figure S7:**
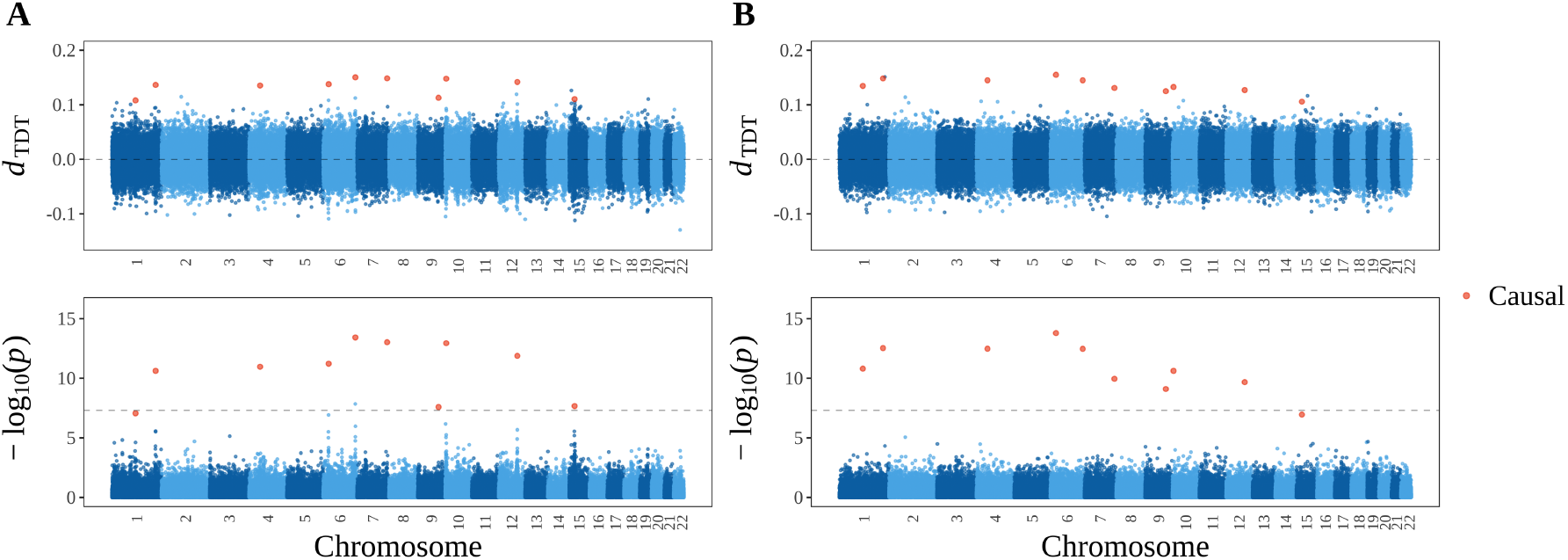
The genome-wide TDT profile for 2,500 affected-only trios. **(A)** Randomization linkage scenario. **(B)** Independent randomization scenario, where the genotypes from (A) were independently permuted to remove the randomization linkage. Genotypes of 5,000 trios are simulated by Section B.5. For both scenarios, the top 2,500 children with the highest value of *ι* + (∑_*i∈*𝒞_ *bG*_*ij*_) + *ϵ*_*j*_ are assigned as affected, setting the trait *Y*_*j*_ = 1 including in the TDT analysis. We draw *ϵ*_*j*_ from Normal(0, 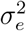), set *ι* = 100, 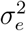 = 1 and determine *b* such that 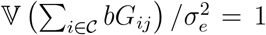. The TDT profile is presented as *d*_TDT_ and log_10_(*p*) at 100,000 SNPs across 22 chromosomes simulated by msprime. The gray dashed line in the bottom panel is at *p*-value = 5 × 10^*−*8^, which is commonly used as a *p*-value threshold in GWAS.

